# Sphingomyelins in mosquito saliva modify the host lipidome to enhance transmission of flaviviruses by promoting viral protein levels

**DOI:** 10.1101/2024.06.14.599058

**Authors:** Hacène Medkour, Lauryne Pruvost, Elliott Miot, Xiaoqian Gong, Virginie Vaissayre, Pascal Boutinaud, Justine Revel, Atitaya Hitakarun, Wannapa Sornjai, Jim Zoladek, R. Duncan Smith, Sébastien Nisole, Esther Nolte-‘t Hoen, Justine Bertrand-Michel, Dorothée Missé, Guillaume Marti, Julien Pompon

**Affiliations:** MIVEGEC, Univ. Montpellier, IRD, CNRS, Montpellier, France; Department of Biomolecular Health Sciences, Faculty of Veterinary Medicine, Utrecht University, Utrecht, The Netherlands; DIADE, Univ. Montpellier, CIRAD, IRD, Montpellier, France; Institute of Molecular Biosciences, Mahidol University, Thailand; Institut de Recherche en Infectiologie de Montpellier (IRIM), Univ. Montpellier, CNRS, Montpellier, France; I2MC, Université de Toulouse, Inserm, Université Toulouse III – Paul Sabatier (UPS), Toulouse, France; MetaboHUB-MetaToul, National Infrastructure of Metabolomics and Fluxomics, Toulouse, France; Laboratoire de Recherche en Sciences Végétales, Metatoul-AgromiX Platform, Université de Toulouse, CNRS, INP, 24 Chemin de Borde Rouge, Auzeville, 31320, Auzeville-Tolosane, France

## Abstract

Mosquito saliva plays a determining role in flavivirus transmission. Here, we discover and elucidate how salivary lipids enhance transmission. Building upon our discovery of salivary extracellular vesicles (EV), we determined that lipids within mosquito EVs, and neither within human EVs nor virions, enhance infection for flaviviruses in primary cell types relevant for transmission. Mechanistically, mosquito EV-lipids specifically promote viral protein levels by reducing ER-associated degradation. Infection enhancement is caused by sphingomyelins within mosquito salivary EVs that elevate sphingomyelin concentration within host cells. Transmission assays showed that mosquito EV-lipids exacerbate disease severity. Our study reveals that EV-associated sphingomyelins within mosquito saliva enhance transmission for multiple flaviviruses by reconfiguring the host lipidome to promote viral protein levels and the resulting skin infection. Our findings open a new dimension centered on lipids in the interplay between hosts, mosquitoes and flaviviruses that determine transmission, unveiling lipids as a new pan-flavivirus target.

**Highlights:**

- Lipids within mosquito extracellular vesicles (EVs) enhance infection in primary skin and immune cells for multiple flaviviruses.
- Mosquito EV-lipids increase flaviviral protein levels by dampening ER-associated degradation.
- Sphingomyelins within salivary EVs are responsible for the infection enhancement by altering host lipidome.
- Co-injection of mosquito EV-lipids exacerbate disease severity.

## Introduction

Several flaviviruses that cumulatively infect half a billion people, cause 250,000 fatalities and €9 billion economic loss annually, are transmitted within mosquito saliva during biting ^1^. Dengue (DENV), Zika (ZIKV) and West Nile (WNV) viruses are the most prevalent flaviviruses and together threaten nearly the entire human population due to the wide geographic distributions of their mosquito vectors ^2–5^. Upon infectious bite, viral multiplication in the skin is required for transmission ^6,7^ and therefore represents a crucial bottleneck to be characterized for the identification of novel targets for much-needed anti-flaviviral interventions ^1^.

Multiple lines of evidence indicate that components in mosquito saliva enhance flavivirus infection ^8–10^ in human cells ^11–13^, mouse models ^6,14–19^, and non-human primates ^20^. Salivary proteins enhancing transmission have been identified. AgBR1 and Nest1 promote inflammation ^21,22^, while sialokinin permeabilizes the endothelial barrier ^23^ - all three proteins amplifying the recruitment of virus-permissive myeloid cells to the bite site. Mosquito LTRIN interferes with the host lymphotoxin-β receptor to inhibit anti-viral NF-κB immune signalling ^24^, the 34-kDa protein reduces type I IFN response ^25^ and salivary AaVA-1 activates pro-viral autophagy ^26^. Recently, we added viral RNA as a new category of salivary transmission-enhancing components ^27^. DENV secrete a subgenomic flaviviral RNA fragment (sfRNA) with anti-immune properties ^28^ in mosquito saliva to inhibit the early cutaneous innate immune response, thereby favouring transmission. Importantly, expectoration of sfRNA occurs within salivary extracellular vesicles (EVs), which we visualized through microscopy ^27^. EVs are spheroid structures delimited by a lipid bilayer membrane and act as cell-free intercellular delivery vehicles, transferring cargo and membrane components such as lipids into adjacent recipient cells ^29^.

Flaviviruses are enveloped single-stranded positive-sense RNA viruses that rely on the host cell lipidome throughout their multiplication cycle ^30–34^. Viral attachment and internalization are mediated by interactions with lipids in the plasma membrane ^35,36^. Translation of the single open-reading frame into a transmembrane polyprotein takes place in endoplasmic reticulum (ER)-associated ribosomes ^37^. Driven by viral non-structural (NS) proteins, ER membranes undergo important structural and compositional rearrangements, inducing invagination to create vesicles that house replication complexes ^38,31^. The negative-sense RNA genome [(-)gRNA] is first synthetized and serves as a template for replication of positive-sense RNA genomes [(+)gRNA]. The resulting (+)gRNA is assembled at the ER but at a different site than replication into a virion enveloped composed of viral structural proteins and ER-derived lipids ^39^. The virion finally egresses through the trans-Golgi secretion network while undergoing maturation, before extracellular release through vacuole fusion with the plasma membrane ^40^.

Concentration of lipids from various classes have to be altered to accommodate flavivirus multiplication cycle. Indeed, classes of structural lipids have specific spatial hindrance and electrostatics which define the biochemical and biophysical properties of infection-induced membrane rearrangements ^34^. Of particular interest, sphingolipids (SLs) are major structural lipids and were regulated in different human cells upon flavivirus infections ^41–44^ and in dengue patient sera ^45^. SLs have a C_18_ sphingosine backbone with a polar headgroup that can be linked to a variety of molecules, producing a range of SLs from the simplest ceramide to the more complex glycoSLs ^46^. When linked to a phosphorylcholine group, ceramides result in sphingomyelins (SM), which are the most abundant SLs in mammalian cells. SMs promote infection for WNV in cells, such as fibroblasts, and in mice ^47,48^, and for Japanese encephalitis virus, another medically-relevant flavivirus, in mouse cells ^49^. Interestingly, extracellular input of SM through cell media supplementation promotes WNV infection ^48^.

In this study, we intertwine the field of flaviviral transmission with lipidomics to discover and reveal how SM lipids contained within mosquito salivary EVs enhance flavivirus transmission. This investigation unveils the mechanistic underpinning of a new category of transmission-enhancing components in mosquito saliva, adding another dimension centered on lipids in the triangular interactions between viruses, mosquitoes and host that determine flavivirus transmission.

## Results

### Mosquito EV-lipids, and neither human EV-lipids nor virion lipids, enhance infection for multiple flaviviruses in transmission-relevant mammalian cell lines

EVs secreted in mosquito saliva can carry their cargo and structural components such as proteins and lipids into vertebrate cells at the bite site ^27,50,51^. To investigate the impact of mosquito EVs on flavivirus infection, we isolated and concentrated EVs from mosquito cell supernatant by ultracentrifugation (Figure 1A), and first supplemented media of permissive human hepatocyte cells with EVs during DENV infection. Supplementation with 0.1 and 1 µl of EV concentrate, where 1 µl corresponded to EVs secreted by ≈ 120,000 mosquito cells over 48h and contained 4.85 µg of proteins, increased intracellular infection as measured by DENV gRNA copies (Figure 1B), while not affecting human cell viability as measured by house-keeping gene levels (Figure S1). Second, we evaluated whether mosquito EVs were internalized by human cells. Concentrated EVs were labelled with a lipid dye and purified through density gradient to remove unbound dye ^52^. Hepatocytes were then exposed to labelled EVs at 37°C before quantifying the proportion of cells that contained the dye and therefore internalized EVs, using high resolution flowcytometry ^53,54^. We observed an increasing proportion of cells containing the EV label from 5% at 2.5h post exposition, 8% at 5h up to 50% at 24h, whereas control cells exposed to material prepared from equal volumes of unconditioned culture medium and subjected to the same staining and purification steps had barely detectable level of labelled cells (Figure 1C). To assess whether mosquito EVs were internalized through endocytosis, we repeated the EV internalization assay maintaining hepatocyte cells at 4°C to inhibit endocytosis. Providing further support for EV internalization, cells maintained at 4°C upon EV exposition did not contain more label than cells exposed to the dye control sample (Figure S2). These initial results suggest that mosquito EVs increase infection by transferring EV components.

**Figure 1.**
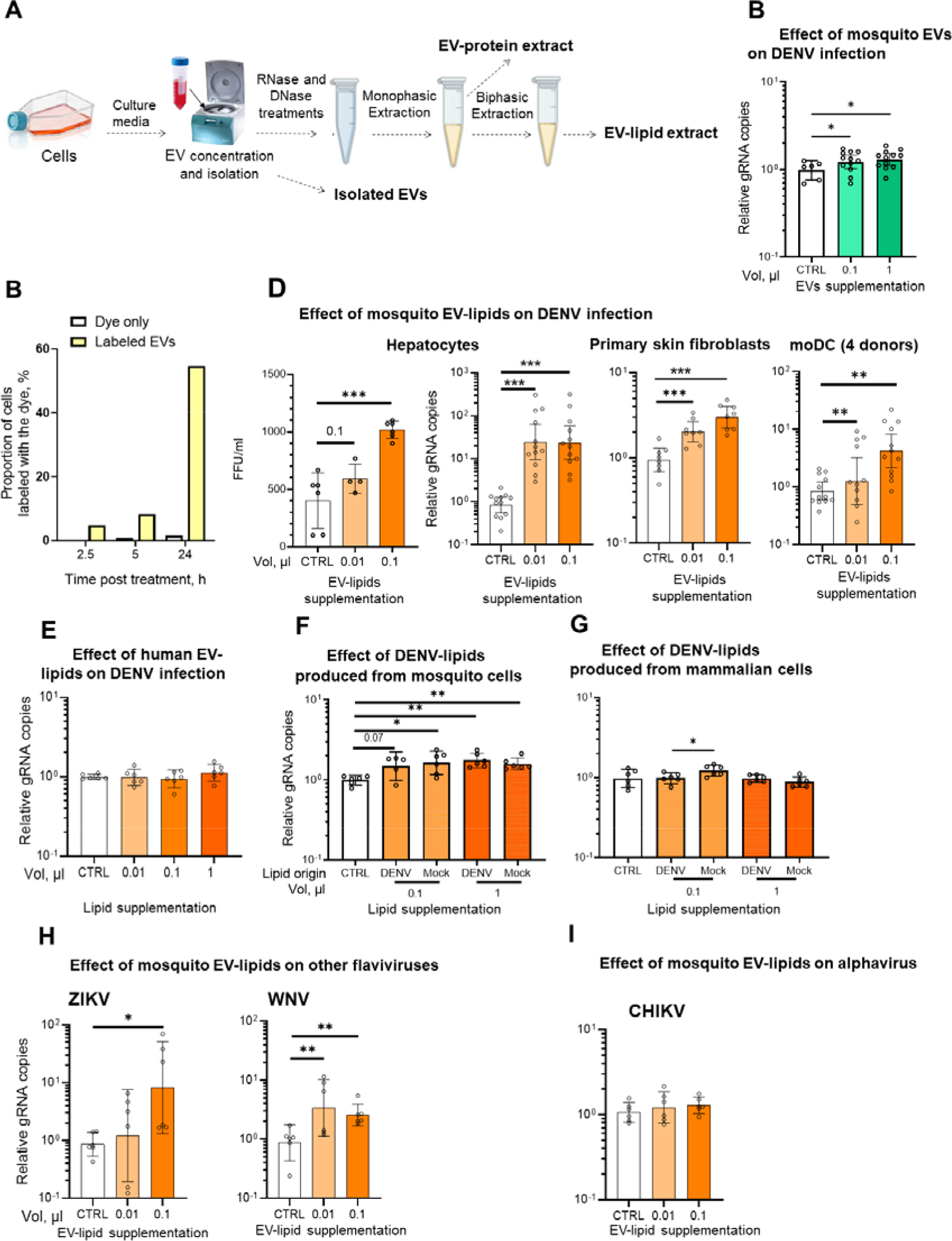
Lipids from mosquito EVs enhance flavivirus infection in multiple human cell types. (A) Scheme of the protocol for EV isolation and extraction of EV-proteins and EV-lipids. (B) Human hepatocytes (Huh7) were infected with DENV at MOI 0.1 upon supplementation with 0.1 or 1 µl of mosquito EV concentrate. (C) Huh7 cells were exposed to mosquito EVs labelled with PKH26 lipid dye at 37°C and dye internalization was quantified 2.5, 5 and 24h later. As controls, cells were exposed to dye-labeled material from an equal volume of similarly processed non-conditioned medium. (D) Huh7, primary neonatal human skin fibroblasts (NHDF) and monocyte derived dendritic cells (moDC) were infected with DENV at MOI 0.1 upon supplementation with 0.01 or 0.1 µl of mosquito EV-lipid extract. EV-lipid ID: LIP 1.1 as in Table S2. (E) Huh7 cells were infected with DENV at MOI 0.1 upon supplementation with 0.01, 0.1 or 1 µl of human EV-lipid extracts. ID of EV-lipid extracts that were used: hLIP 1.1 as in Table S2. (F, G) Huh7 cells were infected with DENV at MOI 0.1 upon supplementation with 0.1 or 1 µl of DENV-lipid extracts produced from mosquito (F) or mammalian (G) cells. Mock indicates supplementation with lipid extracts from the same DENV density fraction from mosquito (F) and mammalian (G) mock-infected cells. Lipid extracts are detailed in Table S3. (H, I) Huh7 cells were infected at MOI 0.1 with two flaviviruses [i.e., ZIKV (H) and WNV (H)] or an alphavirus [i.e., CHIKV (I)] upon supplementation with 0.01 or 0.1 µl of mosquito EV-lipid extract. ID of EV-lipid extracts that were used: LIP 2.2 as in Table S2. (B, D-I) FFU in supernatant and intracellular gRNA were quantified at 72 hpi for DENV, ZIKV and WNV and at 48 hpi for CHIKV. Bars show geometric means ± 95% C.I. from at least 6 biological replicates collected from multiple experiments. Points represent biological repeats. FFU, Focus Forming Unit. CTRL, supplementation with PBS (B), with DMSO (D-I). *, p < 0.05; **, p < 0.01; ***, p < 0.001 as determined by T-test. *See also* Figures S1, S2, S3 and S4; Tables S1, S2 and S3

To disentangle the effects of proteins and lipids contained in mosquito EVs, we extracted EV-proteins and EV-lipids using a modified Bligh and Dyer protocol combined with DNAse and RNAse treatments ^55^ (Figure 1A). The protocol successfully eliminated DNA, RNA, and separated proteins from lipid extracts (Table S1, S2), all of which can influence infection output ^9,10^. To test the effect of EV-proteins, we supplemented hepatocyte cells with 0.051, 0.485, 0.51, or 4.85 µg of EV-protein extracts, encompassing the amount of proteins supplemented through intact EVs. While cell survival was marginally reduced for the higher EV-protein quantities (Figure S3A), none of the EV-protein quantities altered DENV gRNA levels (Figure S3B). In contrast, hepatocyte supplementation with 0.01 or 0.1 µl of EV-lipid extract added to virus inoculum, where 0.1 µl corresponded to lipids extracted from EVs collected from as little as ≈ 6,000 mosquito cells over 48h (Table S2), increased DENV infectious particles in supernatant and intracellular DENV gRNA (Figure 1D), and did not alter cell survival (Figure S4A).

To assess the impact of mosquito EV-lipids on cell types relevant for transmission, we infected primary skin dermal fibroblasts (the most prevalent dermal cell type) and monocyte-derived dendritic cells (moDC) (cutaneous dendritic cells are the primary targets of DENV following skin inoculation ^6,56^) upon mosquito EV-lipid supplementation. In both primary cell types, we observed an increased infection (Figure 1D), while cell viability was not affected (Figure S4B-C). These results reveal that lipids contained in mosquito EVs enhance DENV infection in multiple transmission-relevant cell types.

Since all cells produce EVs ^29^, we next evaluated the impact of EV-lipids from human cells by similarly extracting lipids from hepatocyte EVs and supplementing human cells during DENV infection with 0.01, 0.1 or 1 µl of human EV-lipids, where 1 µl corresponded to lipids extracted from EVs collected from ≈ 65,000 human cells over 48h (Figure 1A; Table S2). In support of a specific function for mosquito EV-lipids, human EV-lipids did not influence DENV infection (Figure 1E), nor cell viability (Figure S4D).

DENV envelop contains lipids, which composition partially varies when viruses derive from mosquito or mammalian cells ^57,58^. To evaluate the impact of DENV-lipids on DENV infection, we produced DENV in mosquito and monkey cells, and isolated virions from EVs ^59^ using discontinuous sucrose density gradient. The purified DENV samples were obtained from our previously-published study, which reported an absence of EV markers (i.e., Alix) in the DENV density fraction^58^, suggesting at least partial separation of DENV from Alix-containing EVs. We confirmed DENV isolation by detecting high quantity of DENV gRNA (Table S3). We then extracted lipids from the DENV density fractions from either mosquito or mammalian cells and, as controls, extracted lipids from the same density fractions from mock-infected samples (Table S3). Using hepatocyte cells, we observed that infection was increased upon supplementation with lipids from DENV produced in mosquito cells (Figure 1F). However, infection was similarly enhanced when cells were supplemented with lipids from mock-infected mosquito cell supernatant. Since EVs and DENV have partially overlapping density ^59,60^, the mock-infected fraction probably contained EVs, which did not possess or had undetectable amount of the EV marker ^58^ and the increased infection upon supplementation with mock-infected lipids was likely caused by residual mosquito EV-lipids. In contrast, supplementation with either DENV or mock-infected lipids from mammalian cells did not alter DENV infection (Figure 1G). Cell viability was unaffected in all conditions (Figure S4E, F). These results indicate the lack of impact on infection for lipids contained within virions.

Finally, we examined the impact of mosquito EV-lipids on infection for other arboviruses. Both flavivirus WNV and ZIKV infections were increased by mosquito EV-lipid supplementation (Figure 1H; Figure S4G,H). Contrarily, replication of chikungunya virus (CHIKV), the most prevalent mosquito-borne alphavirus ^61^, was not influenced by mosquito EV-lipids (Figure 1I; Figure S4I). Altogether, our results reveal that lipids contained within mosquito EVs - but neither within human EVs nor flavivirus envelop - enhance infection for multiple flaviviruses - but not alphaviruses - in multiple transmission-relevant cell types.

### Mosquito EV-lipids specifically promote viral translation by dampening ER-associated degradation

To determine how mosquito EV-lipids enhance flaviviral infections, we assessed the impact of mosquito EV-lipids on each stage of the DENV cellular cycle. At the onset of infection, attachment and internalization, as quantified by numbers of attached and internalized (+)gRNA, were not affected (Figure 2A-B). In contrast, early translation, as estimated by virus protein quantity from 3-6 hours-post infection (hpi), was progressively enhanced by mosquito EV-lipids with a significant increase at 6 hpi (Figure 2C). Replication was estimated as the kinetics of antigenome [(-)gRNA] production from 1-24 hpi. While (-)gRNA was not detected at 1 hpi as expected at the onset of the cycle, (-)gRNA was detected from 3 hpi onward but its levels were not affected by mosquito EV-lipids up to 18 hpi (Figure 2D). At 24hpi, however, we observed an increase in (-)gRNA upon mosquito EV-lipid supplementation, which is consistent with an enhanced replication caused by higher quantities of viral proteins^32^. Finally, virion assembly and excretion were evaluated by quantifying infectivity of secreted particles as the ratio of (+)gRNA:FFU in supernatant. Although mosquito EV-lipids enhanced the production of infectious particles (Figure 1D), virion infectivity was not altered (Figure 2E). Together, these results suggest that mosquito EV-lipids specifically promote viral translation, resulting in an increased amount viral proteins.

**Figure 2.**
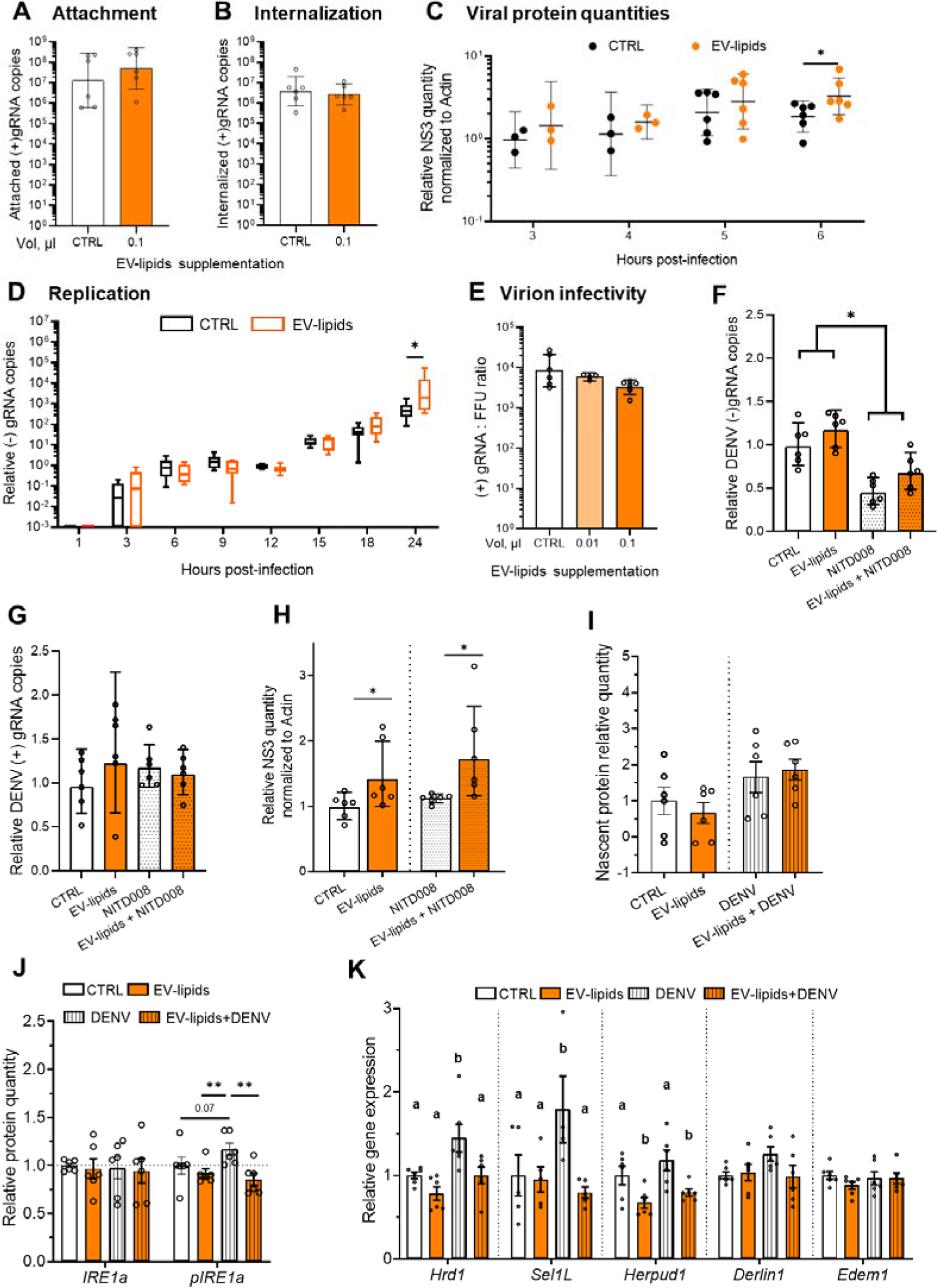
Mosquito EV-lipids increase viral protein levels by altering ER-associated degradation. (A-E) Huh7 cells were infected with DENV at MOI 0.1 upon supplementation with 0.1 µl of mosquito EV-lipids or DMSO (CTRL) before quantifying attached (A) and internalized viruses (B), viral translation as early production of NS3 protein (C), viral replication as copies of (-)gRNA (D), and viral infectivity as the (+)gRNA:FFU ratio in supernatant (E). ID of EV-lipid extracts that were used: LIP 1.1 and LIP 2.2 as in Table S2. (F-H) Huh7 were treated with an inhibitor of viral genome replication (NITD008) before and during infection with DENV at MOI 0.1 upon supplementation with 0.1 µl of mosquito EV-lipids or DMSO (CTRL). Internalized (+)gRNA (F), viral replication as copies of (-)gRNA copies (G), and viral translation as early production of NS3 protein (H) were quantified at 6 hpi. ID of EV-lipid extracts that were used: LIP 2.5_2 and LIP 2.5_3 as in Table S2. (I-J) Huh7 cells were infected with DENV at MOI 0.1 upon supplementation with 0.1 µl of mosquito EV-lipids or DMSO (CTRL) before quantifying nascent protein production (I), IRE1a and pIRE1a proteins (J) and expression of ER-associated degradation genes (K) at 6 hpi. ID of EV-lipid extracts that were used: LIP 7.1 as in Table S2. (A-C, E-H, J) Bars and lines show geometric means ± 95% C.I. Points represent biological repeats collected from multiple experiments. (D) Tukey box and whiskers from six biological replicates collected from two independent experiments. (I, J) Bars show means ± SEM. Points represent biological repeats. (C, D) *, p < 0.05 as determined by one-tailed t-test. (F, H, J) *, p < 0.05; **, p < 0.01 as determined by one-tailed t-test. (K) Different letters indicate significant differences, as determined by LSD’s test (p < 0.05). *See also* **Figure S5**

To confirm that mosquito EV-lipids promote viral translation, we inhibited viral replication by supplementing the cell media prior and during the course of infection with NITD008, a potent viral RNA synthesis inhibitor ^62^, and quantified viral translation at 6 hpi. First, at 6 hpi, we confirmed a drastic, although not complete, inhibition of replication by quantifying a 66% reduction in (-)gRNA levels upon NITD008 treatment – an inhibition that was maintained when mosquito EV-lipids were supplemented (Figure 2F). Second, at 6 hpi, we monitored intracellular (+)gRNA quantities, which, in absence of viral replication, originates from internalised virions. Confirming that mosquito EV-lipids do not influence attachment and internalisation, (+)gRNA quantities were unaltered by mosquito EV-lipids, NITD008 and their combination (Figure 2G). Eventually, we observed that mosquito EV-lipids promote viral translation, even when viral replication is inhibited by concomitant NITD008 treatment, by observing a 66% increase in viral protein quantity – an increase that is comparable to that observed without NIT0008 treatment (Figure 2H). Therfore, chemical inhibition of viral replication further supports that mosquito EV-lipids increase flaviviral infection by enhancing viral protein quantity.

We then elucidated the mechanism by which EV-lipids increase viral protein levels. First, we rejected the hypothesis that EV-lipids induce an overall translation increase by showing that neither EV-lipids nor EV-lipids in combination with DENV infection enhanced nascent protein quantity (Figure 2I), as previously reported for DENV infection ^63,64^. Second, because the unfolded-protein response (UPR) is triggered by infection-induced ER stress ^65,66^, we tested whether EV-lipids regulated the UPR by quantifying the phosphorylation status of the signal activator IRE1a at 6 hpi ^67,68^, when viral protein levels are heightened (Figure 2C). While total amount of IRE1a was not influenced by either DENV infection, EV-lipids or infection upon EV-lipid supplementation, we observed a moderate increase in the amount of phosphorylated/activated IRE1a (pIRE1a) upon DENV infection (Figure 2J), confirming the previously-reported marginal induction of the UPR upon flavivirus infection ^37,64,69^. Strikingly, however, EV-lipid supplementation alone or in combination with DENV infection maintained pIRE1a levels lower than those upon infection (Figure 2J). Third, we determined whether EV-lipids influence the UPR-induced ER-associated degradation (ERAD), which function is to restore ER homeostasis by transporting misfolded proteins to the proteasome for degradation ^70,71^. ERAD activation results in transcription of multiple genes, such as *Hrd1*, *Sel1L*, *Herpud1*, *Derlin1* and *Edem1*, which are involved in the different branches of the ERAD pathway ^72^. By measuring expression of each of the above genes, we observed 3 groups of regulation patterns; *Hrd1* and *Sel1L* were induced by DENV infection, whereas their infection-induced activation was inhibited upon EV-lipid supplementation; *Herpud1* was not induced by infection but was downregulated by EV-lipids alone or in combination with infection; and *Derlin1* and *Edem1* were neither regulated by EV-lipids, infection nor the combination of infection and EV-lipids (Figure 2K). Altogether, our results strongly suggest that EV-lipids dampen infection-induced UPR, resulting in a lower activation of some branches of the ERAD pathway to diminish viral protein degradation and, thereby, enhancing viral protein quantity.

Since innate immune response potently restricts flaviviral infections ^73^, we also evaluated the impact of mosquito EV-lipid supplementation on *IFN-β* and two interferon-stimulated genes (ISG) (i.e., *MX1* and *CXCL10*). At both 24- and 72-hours post-treatment, none of the immune-related genes were regulated by mosquito EV-lipids, whereas these genes were clearly induced by DENV infection (Figure S5A-C). These results suggest that the mosquito EV-lipid infection enhancement is not caused by immune inhibition.

### Sphingomyelins within mosquito salivary EVs are responsible for the infection enhancement

To identify the lipid class responsible for the mosquito EV infection-enhancement, we first described the composition of mosquito EV-lipids using targeted lipidomics to obtain quantitatively comparable results across the different classes of lipids ^74^. Performing specific quantifications for SL, phospholipids, neutral lipids and fatty acids, we detected and quantified 107 lipid species (Dataset S1). The most abundant lipids were neutral lipids and phospholipids such as PC and PE (Figure 3A; Dataset S1). Although EV-lipid composition varies with cell types, our first-ever description of mosquito EV-lipids broadly corresponds to the lipid composition of mammalian EVs^51^. Of particular interest, with our method, the 11 SM species represented 1.34% of the total lipid mass from mosquito EV-lipids (Dataset S1).

**Figure 3.**
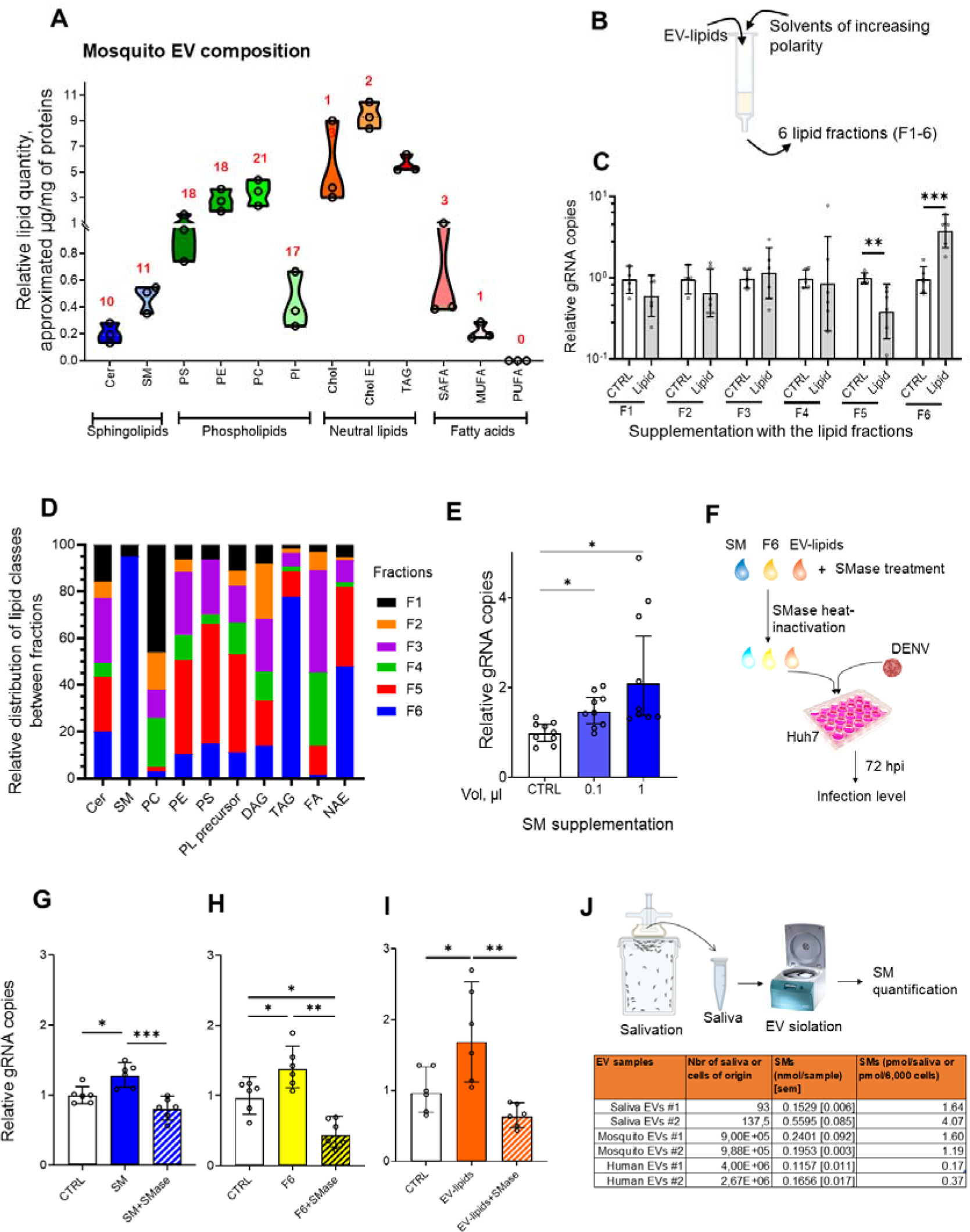
Sphingomyelins (SM) in mosquito EVs are responsible for the infection increase and are detected in mosquito salivary EVs. (A) Quantitative targeted lipidomics on mosquito EV-lipid extracts. Data derived from three biological replicates, represented by different circles. Red numbers indicate number of detected lipid species within each lipid class. (B) Scheme of the fractionation protocol for mosquito EV-lipids into 6 fractions through solid-phase extraction (SPE). (C) Huh7 cells were infected with DENV at MOI of 0.1 upon supplementation with 0.1 µl of the different lipid fractions (F1-6) or corresponding control (CTRL). (D) Lipid class relative composition of the mosquito EV-lipid fractions by untargeted lipidomics. Total of relative proportions within each lipid class equals 100%. F1-6, SPE fractions 1-6. (E) Huh7 cells were infected with DENV at MOI 0.1 upon supplementation with 0.1 or 1 µl of commercially-available pig brain SM. CTRL, supplementation with 1 µl of H_2_O. (F) Experimental design for SMase pre-treatment of lipids prior supplementation. (G-I) Huh7 cells were infected with DENV at MOI 0.1 upon supplementation with 1 µl of SM (G), 0.1 µl of F6 (H) or 0.1 µl of EV-lipids that had been pre-treated with SMase and heat-inactivated. In absence of SMase treatment, lipids were similarly heated. CTRL, supplementation with DMSO including heat-inactivated SMase. ID of EV-lipid extracts that were used: LIP 2.5_3 as in Table S2. (J) Quantification of SM in EVs from mosquito saliva. Two saliva pools were quantified. Averages of SM quantify from 2 replicates of mosquito and human cell EVs are shown. (A, D) Cer, Ceramide; SM, Sphingomyelin; PS, Phosphatidylserine; PC, Phosphatidylcholine; PE, Phosphatidylethanolamine; PI, Phosphatidylinositol; Chol, Cholesterol; Chol E, Cholesterol esters; SAFA, Saturated fatty acids; MUFA, monounsaturated fatty acid; PUFA, polyunsaturated fatty acid; TAG, triacylglycerides; DAG, diacylglycerides; FA, Fatty acids; NAE, N-acylethanolamine. (C, E, G-I) Intracellular gRNA copies were quantified at 72hpi. Bars show geometric means ± 95% C.I. Points represent biological repeat collected from three (E) or two (C, G-I, K) different experiments. *, p < 0.05; **, p < 0.01; ***, p < 0.001 as determined by one-tailed T-test. *See also* **Figure S6, Dataset S1, S2**

Second, we fractioned mosquito EV-lipids according to lipid polarity using solid-phase extraction (SPE) columns and sequential elutions with solvents of increasing polarity (Figure 3B). The resulting 6 fractions and the corresponding controls (CTRL), obtained by eluting empty SPE column with the corresponding solvents, were subjected to lipid extraction (Figure 1A). Human cells were then infected with DENV upon supplementation with fractioned lipid extracts and infection was quantified at 72 hpi. While cell viability was not altered in any conditions (Figure S6A, B), fractions 1-4 did not affect infection and fraction 5 reduced infection levels (Figure 3C). Strikingly, fraction 6 increased infection and therefore contained the infection-enhancing lipids. We then identified lipids through untargeted global lipidomics, which enable agnostic lipid identification but only permit within-lipid class relative quantification ^74^. We detected 228 lipid species, mostly neutral lipids such as diacylglycerides (DAG) and triacylglycerides (TAG), and phospholipids such as PC and PE (Dataset S2), in coherence with our targeted lipidomics description (Dataset S1). As compared to the other fractions, fraction 6 was enriched in SM, containing 95% of total SM (Figure 3D; Dataset S2).

Third, we evaluated the function of SMs contained in mosquito EVs. While supplementing human cells with commercially-purified SMs increased DENV infection (Figure 3E), as previously shown for WNV ^48^, we tested whether SMs within mosquito EVs were responsible for the infection enhancement. To this end, we removed SMs from mosquito EV-lipid extracts through a sphingomyelinase (SMase) treatment and supplemented cells with the resulting SM-free lipids during DENV infection (Figure 3F). We validated our SM removal approach by showing that SMase treatment abrogated the infection enhancement caused by purified SMs (Figure 3G). We then observed that the infection enhancement was similarly abrogated when fraction 6 (Figure 3H) and EV-lipid extracts (Figure 3I) were pre-treated with SMase. Our results decisively establish that SMs in mosquito EVs are responsible for the enhancement of flaviviral infections.

Finally, to translate our findings to real-world transmission, we quantified SMs in mosquito salivary EVs. Since saliva minute volumes do not permit the exhaustive biological and biochemical characterization we conducted with mosquito cell-derived EVs, we collected saliva from 93 and 137 mosquitoes, enriched EVs and used a colorimetric kit to quantify SMs (Figure 3J). Our samples contained 1.64 and 4.07 pmol of SMs per saliva (Figure 3J). In comparison, mosquito EVs used to produce 0.1 µl of EV-lipid extracts, which increased flavivirus infection (Figure 1) contained an average of 1.39 pmol of SMs (Figure 3J). Interestingly, human EVs used to produce 0.1 µl of EV-lipid extracts, which had no impact on infection (Figure 1D), only contained an average of 0.27 pmol of SMs, indicating that EVs derived from the human cells we used have less infection-enhancing SMs (Figure 3J). Altogether, our investigation demonstrates that SMs found in EVs from mosquito cells and mosquito saliva enhance flavivirus infection.

### Sphigomyelins within mosquito EVs amplify infection-driven increase in sphingomyelin concentration in host cells

To pinpoint the cellular lipidome associated with the EV-lipid-induced enhancement of flaviviral infection, we described how mosquito EV-lipids reconfigure the host cell lipidome. For this, we conducted global untargeted lipidomics on human cells treated with: (i) neither EV-lipids nor DENV, (ii) mosquito EV-lipids, (iii) DENV, and (iv) DENV with mosquito EV-lipids. Reasoning that different quantities of viral proteins may alter the cell lipidome, we collected cells at 4 hours post-treatment (hpt), before the increase in viral protein levels (Figure 2C). We detected 280 lipid species, spanning different lipid classes including SL, phospholipids, neutral lipids and fatty acids (Dataset S3). In spite of the early time point (i.e., 4 hpt), 80 lipids (28.5% of the total) were significantly regulated in at least one condition (Dataset S3). We clustered the regulated lipids according to their expression patterns, revealing 9 regulation clusters across the four conditions (Figure 4A). To decode this complex dataset, we aligned with previous studies showing that flaviviruses modulate cellular lipidome for their benefits ^75–77^ and posited that infection-induced lipid regulation indicated a pro-viral environment. To simplify interpretation, we plotted the trends in regulation clusters in a heatmap (Figure 4B).

**Figure 4.**
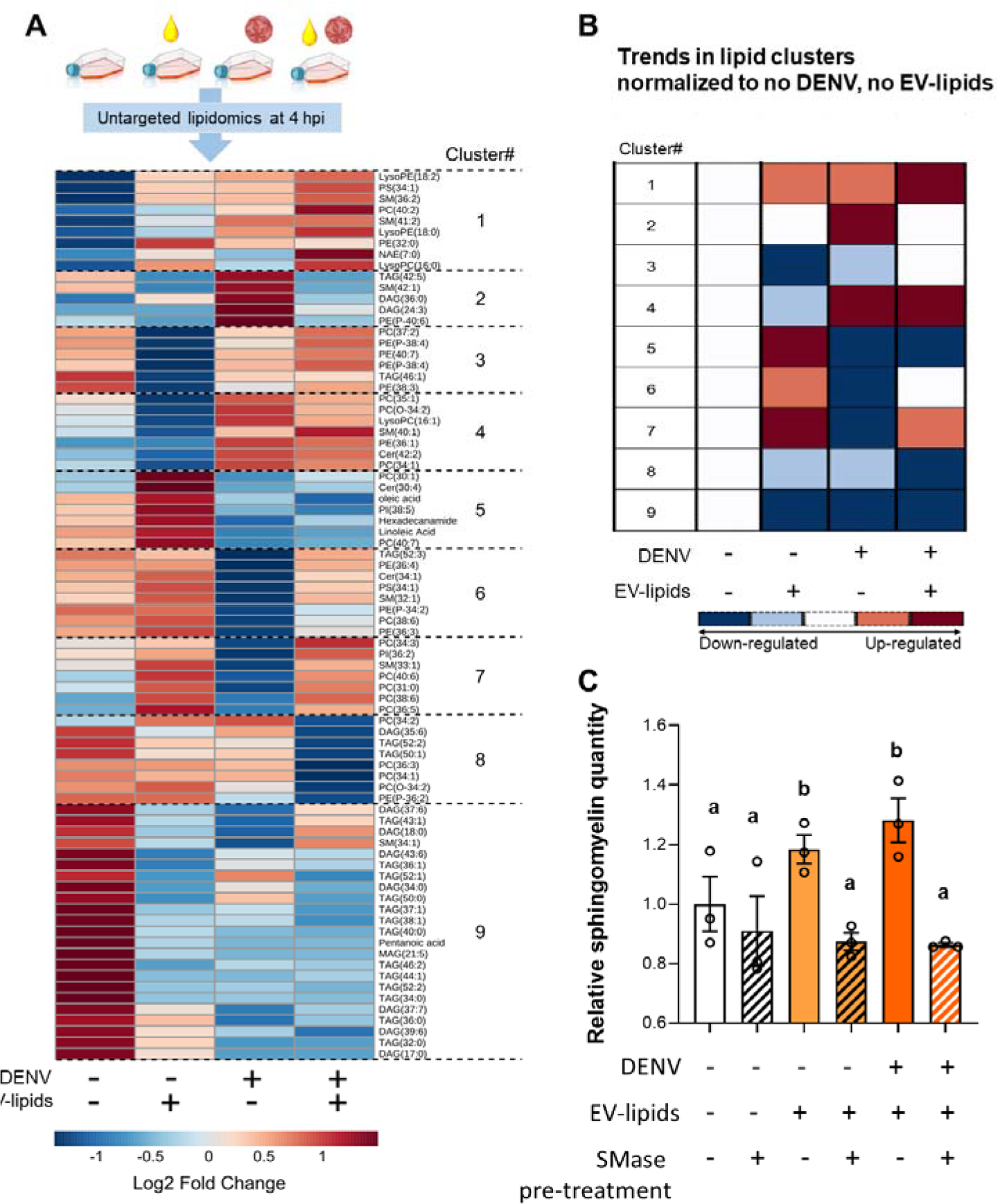
Mosquito EV-lipids reshape human cellular lipidome, partially amplifying DENV-induced reconfiguration. (A) Regulation of cellular lipids. Huh7 cells were inoculated with 0.1 µl of mosquito EV-lipids, DENV at MOI 0.1 or both EV-lipids and DENV. Control cells were neither supplemented with lipids nor infected. Untargeted lipidomics of cells was conducted after 4 h. ID of EV-lipid extracts that were used: LIP 2.2 as in Table S2. Fold changes and clustering of significantly regulated (|log2 fold change| > 1; p-value < 0.05) lipids. Values were averaged from three biological replicates. (B) Schematic representation of the regulation of the 9 lipid clusters. Changes were normalized to no-DENV, no-lipid condition. (C) Regulation of cellular sphingomyelin by DENV and EV-lipids upon SMase pre-treatment at 4h post exposure. Different letters indicate statistical differences (p < 0.05). *See also* **Dataset S3**

Cluster 1 comprises 9 lipids that were upregulated by infection and by mosquito EV-lipids (Figure 4A, B). Notably, their upregulation was amplified when infection was supplemented with mosquito EV-lipids. Based on our postulate that infection-modulated lipids create a conducive environment for viral proliferation, lipids of cluster 1 were identified as pro-viral. Cluster 1 lipids included 2 SMs, which promote WNV infection *in vivo* ^47,48^, 3 lysophospholipids, known to increase WNV genome replication ^76^, 3 phospholipids that influence DENV genome replication ^75^ and 1 short fatty acid.

Lipids from clusters 2 and 4 were upregulated by infection (Figure 4A, B). However, mosquito EV-lipids did not regulate cluster 2 lipids and downregulated cluster 4 lipids, suggesting that the lipids in clusters 2 and 4 were not associated with a pro-viral environment. Furthermore, concomitant treatment with DENV and EV-lipids abrogated infection-induced regulation of cluster 2 lipids and did not amplify infection-induced regulation of cluster 4 lipids.

Lipids from clusters 3, and 5-9 were downregulated by infection (Figure 4A, B). While mosquito EV-lipids alone similarly downregulated cluster 3, it was surprising to observe that the infection-induced regulation was annihilated when infection was combined with EV-lipids. Lipids from clusters 5-7 were upregulated by mosquito EV-lipids, antagonizing the infection-induced pro-viral regulations. In contrast, lipids from clusters 8 and 9 were downregulated by mosquito EV-lipids as upon infection, potentially creating a pro-viral environment. Furthermore, the combination of EV-lipids and infection amplified the downregulation of the cluster 8 lipids and maintained the infection-induced reduction of the cluster 9 lipids. Cluster 8 mostly comprised phospholipids [4 phosphatidylcholine (PC) and 1 phosphatidylethanolamine (PE)], which are involved in membrane structure ^78^, whereas the 23 lipid species from cluster 9 were almost exclusively fatty acids (TAG and DAG), involved in energy metabolism ^79^. Altogether, we showed that mosquito EV-lipids induce an early and complex modulation of multiple classes and highlighted lipids which infection-induced regulation were amplified by mosquito EV-lipids, potentially creating a pro-viral environment. Of particular interest were the cluster 1 pro-viral SMs, which infection-induced upregulation was amplified by mosquito EV-lipids.

We next tested whether EV-associated SMs were responsible for the increase in SM concentration in host cells. We exposed human cells to DENV and mosquito EV-lipids, which had been depleted or not of SM by a SMase pre-treatment as previously (Figure 3F), and quantified cellular SM concentration at 4 h post-exposition. First, we showed that exposition to heat-inactivated SMase alone did not alter cellular SM concentration as compared to control (compare bars 1 and 2; Figure 4C). Second, we confirmed that EV-lipids elevated cellular SM concentration (compare bars 1 and 3; Figure 4C), and, strikingly, showed that SMase pre-treatment abrogated this increase (compare bars 2 and 3; Figure 4C). Finally, we reproduced the increase in cellular SM concentration observed using global lipidomics when DENV infection was supplemented with EV-lipids (compare bars 1 and 5; Figure 4C), and again showed that this increase was absent with SMase pre-treated EV-lipids (compare bars 5 and 6; Figure 4C). Altogether, these results characterize how EV-lipids quickly reconfigure the host lipidome, demonstrating that SM within EVs are responsible for the increased SM concentration in host cell.

### Mosquito EV-lipids aggravate disease severity in a mouse model of transmission

To evaluate the impact of mosquito EV-lipids on transmission, we inoculated a wild-type mouse model susceptible to WNV infection ^21^ with a sub-lethal dose of WNV in combination with 0.1 or 1 µl of mosquito cell EV-lipids (Figure 5A), covering the range of SM quantities present within one saliva (Figure 3J). Control mice received either DMSO alone (No inf.) or the WNV inoculum mixed with DMSO (CTRL). Inoculum volume was designedly small (i.e., 10 µl) and injected in the dermis to mimic bite delivery of saliva in the skin.

**Figure 5.**
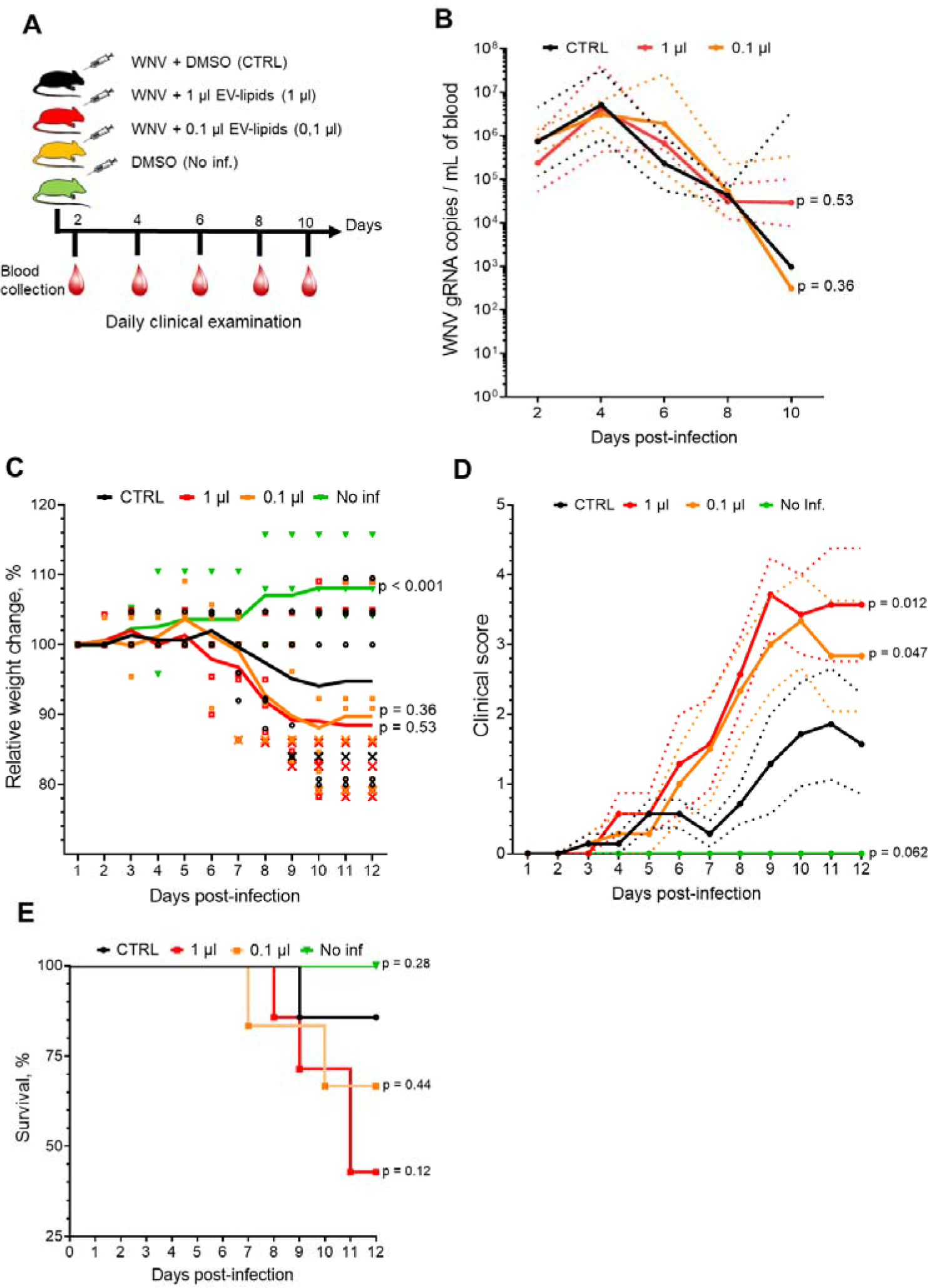
Intradermal co-inoculation of WNV with mosquito EV-lipids increases disease severity in mice. (A) Adult C57BL/6J mice (6- to 8-week-old) were inoculated intradermally with 10^3^ WNV supplemented with 0.1 or 1 µl of mosquito EV-lipids. Control mice (CTRL) were inoculated without lipid supplementation. Non-infected mice (No inf.) were injected with DMSO. Mice were daily monitored for weight loss, clinical signs and survival for 12 days. RNAemia was quantified at 2, 4, 6, 8, and 10 dpi. N, 7 for lipid supplementation and CTRL conditions. N, 6 for no infection condition. ID of EV-lipid extracts that were used: LIP 1.1 and LIP 2.5_1 as in Table S2. (B-E) RNAemia (B), Relative weight change (C), Clinical score (D), and survival (E). (B) Lines show geometric means and dotted lines 95% C.I. (C, D) Lines show average values. (C) Each point represents one mouse. x, stands for weight variation in deceased mice. (D) Dotted lines indicate SEM. (E) Lines indicate survival curves. (B-E) p-values indicate differences with CTRL for the interaction between time and treatment. *See also* **Figure S7; Table S4**

We first monitored RNAemia in blood using absolute quantification (Figure S7) every two days for 10 days. Viremia quantification using titration was not conducted as it is usually undetectable in wild-type mice ^80–82^. In all three infectious conditions, while WNV gRNA was detected as early as 2 days post infection (dpi) with a peak at 4 dpi ranging from 3.1 - 5.2 x 10^6^ gRNA / ml, EV-lipid co-inoculation had no impact (Figure 5B). Although serum RNAemia does not usually reflect disease severity ^80–82^, our results validated active infection in mice. Second, we analyzed mouse weight variations and observed that infected mice lost weight (Figure 5C), as expected from WNV infection ^80–82^. Strikingly, the weight loss was amplified, although not significantly, when WNV was co-inoculated with EV-lipids (Figure 5C). Third, we evaluated symptoms according to a clinical score ranging from 0 – 5 (Table S4), considering key WNV symptoms in mice and where 5 corresponds to death ^83^. In all infectious conditions, clinical signs followed a similar pattern (Figure 5C) – first symptoms on day 2, progressively aggravating until day 9-11, at which time clinical scores plateaued or decreased. Interestingly, as compared to mice solely injected with WNV, symptom severity increased with co-injection of 0.1 µl (Two-way repeated-measured ANOVA; time effect, p-value = < 0.0001; infection effect, p-value = 0.147; interaction, p-value = 0.0471 µl) and 1 µl (time effect, p-value = < 0.0001; infection effect, p-value = 0.0193; interaction, p-value = 0.012) of EV-lipids (Figure 5D). Finally, while infection reduced survival to 85 %, as expected from the sublethal inoculum, co-inoculation with 0.1 and 1 µl of mosquito EV-lipids further diminished survival to 66 and 44%, respectively (Figure 5E). Altogether, by performing *in vivo* infections that mimic bite-initiated transmission, we demonstrated that co-inoculation of mosquito EV-lipids amplifies disease severity in a dose-dependent manner.

## Discussion

While proteins and viral RNA are well-known salivary factors that influence skin infection ^9,26,27^ and the resulting transmission ^8,18,84^, our investigation uncovers lipids as a previously unrecognized category of salivary components that enhance the transmission of multiple flaviviruses. This discovery introduces a novel lipid dimension to the intricate interactions between viruses, hosts and mosquitoes that governs transmission dynamics. Simulated by our initial identification of EVs in mosquito saliva ^27^, we explored the role of mosquito EVs in flavivirus transmission. Our findings reveal that mosquito EVs promote flavivirus infection via the lipids they carry. Furthermore, the EV-lipid-mediated infection-enhancement is conserved across flaviviruses and multiple transmission-relevant cell lines, unveiling a novel pan-flaviviral mechanism of transmission enhancement. Mechanistically, EV-lipids specifically bolster viral protein quantities by mitigating UPR-induced ERAD-mediated protein degradation. Within mosquito EVs, we identified SMs as the key lipid class responsible for the infection enhancement. In the host, EV-lipids induce a complex reconfiguration of the cellular lipidome, in which EV-associated SMs amplify the infection-driven upregulation of pro-viral SMs. Finally, by detecting high concentration of SMs within mosquito salivary EVs and showing that mosquito EV-lipids enhance disease severity in a mouse model of transmission, our study establishes that mosquito salivary EV-lipids increase transmission. Altogether, our study demonstrates that SMs contained in mosquito salivary EVs create a pro-viral lipidome environment in host cells, increasing flaviviral protein levels by dampening protein degradation, thereby enhancing initial skin infection and subsequent transmission for all flaviviruses.

Our findings indicate that mosquito EV-lipids, mostly SMs, mitigate UPR-induced ERAD to increase flaviviral protein quantities and infection. UPR activation by flavivirus infection is well-established ^37,64,69^ and was shown to reduce WNV multiplication ^66^. These prior studies in combination with our data establish UPR activation as an anti-viral mechanism, which inhibition by mosquito EV-lipids promote flaviviral infection. In contrast to flaviviruses, alphaviruses like CHIKV are not susceptible to UPR anti-viral mechanisms as their nsP2 protein inhibits the expression of UPR transcription factors ^85^. Accordingly, we did not report any pro-CHIKV function for mosquito EV-lipids. The UPR activation by misfolded proteins, resulting from infection, induces the phosphorylation of the transcription factor IRE1a^86^. However, the ER transmembrane localisation of IRE1a protein renders IRE1a activation vulnerable to lipid membrane composition, which can alter protein reactivity ^87^. Accordingly, dysregulation of SL biosynthesis induces UPR ^88–90^. We therefore propose that the host cellular SM reconfiguration induced by mosquito EV-associated SMs alters the ER membrane lipid composition, hampering IRE1a activation. Upon phosphorylation IRE1a induces the expression of more than 10 ERAD-related proteins with different functions, resulting in the transport of identified misfolded proteins to the proteasome for degradation. We reported that mosquito EV-lipids inhibit the expression of several ERAD genes, providing a rationale for increased viral protein levels when infection was supplemented with mosquito EV-lipids. Our mechanistic characterization reveals a novel lipid-based mechanism by which flavivirus infection is promoted.

Co-inoculation of viruses with EV-lipids in mice resulted in aggravated disease severity. Upon biting, flaviviruses replicate in various skin cells – prior studies reported infections in fibroblasts ^91,92^, keratinocytes ^11,93^, dermal dendritic cells (DC) ^56,94^, and Langerhans cells (LC) ^95^. A few hours post biting, virus-permissive myeloid cells are recruited to the bite site, amplifying local infection and triggering systemic spread by migrating to lymph nodes ^9,18,23,94^. Our findings revealing that EV-lipid-mediated infection-enhancement is consistent across multiple skin cell types, including primary fibroblasts and myeloid cells, indicate that salivary EV-lipids promote local infection at the bite site. Direct effects of salivary EV components are likely short-lived, as aqueous components delivered in the dermis are absorbed within hours ^96^ and EVs are usually internalized within hours ^97,98^. Our study reports an extensive lipidome reconfiguration as early as 4 hours post exposition to EV-lipids, which is concomitant with a quantifiable increase in viral proteins at 6 hpi. Based on prior studies showing that altering the initial skin infection influences transmission and disease severity in mice ^7^, we posit that SMs from mosquito salivary EVs enhance initial infection in skin cells, heightening transmission and disease severity. The pronounced influence of EV-lipid co-inoculation on disease severity further underscores the critical role of initial cutaneous infection in determining overall disease progression.

In conclusion, our discovery of a pan-flavivirus transmission-enhancing function for mosquito salivary lipids opens new opportunity for much-needed broad-spectrum therapeutics. Existing lipid-altering drugs ^99^ could be leveraged to derail pro-viral lipid reconfiguration at the bite site. More broadly, while the effects of EV-lipids start to garner attention ^51^, our study provides the first evidence that lipids derived from EVs influence viral infection. Overall, our mechanistic insights into the role of salivary lipids in bite-initiated flavivirus infection expand the range of salivary component categories that govern flaviviral transmission.

## Study limitations

We acknowledge several limitations in our study. First, mechanistic characterization was conducted using EVs produced from a cellular model. The mosquito cell line used was derived from larvae and the organ from which it originates is unknown ^100^. However, minute quantities of saliva do not permit lipid extraction in sufficient quantities for the analyses. Although there might be variability in EV lipid composition between cellular model and mosquito saliva, we validated the presence of the functional SMs in EVs from saliva as from mosquito cells. Second, we used lipids extracted from different batches of EVs and, although the infection-enhancement was reproducible, we observed variability in the intensity of infection increases. The variability may stem from differences in cell culture conditions that influence EV lipid composition ^51^. To inform the readers about this potentially confounding effect, we indicated the details of the EV batch that was used in each experiment. Third, in our *in vivo* transmission assay, we mimicked bite-initiated transmission by intradermal inoculation, which may not reproduce the complex process of biting. Altogether, we combined *in vitro* and *in vivo* approaches to support the discovery of salivary lipids as a new category of transmission-enhancing compounds.

## Supporting information

Key Ressources

## Acknowledgements

We extend our gratitude to the members of the Pompon’s Laboratory at MIVEGEC, IRD, for their valuable suggestions and support. Special thanks to Dr. Thierry Durand and Juliette Van-dijk from CNRS, Montpellier for their assistance. Our appreciation goes to Nathalie Barougier and Drs Sylvie Cornelie and Idris Mhaidi for their help in mouse management. Additionally, we are grateful to the VectoPole team in Montpellier, particularly Bethsabée Scheid and Carole Ginibre for providing the BORA eggs. Lipidomic analysis were performed at MetaToul (Toulouse metabolomics & fluxomics facilities, www.mth-metatoul.com), which is part of the French National Infrastructure for Metabolomics and Fluxomics MetaboHUB-ANR-11-INBS-0010. We are grateful to Nancy Geoffre, Amelie Perez, Océane Delos and Anaele Durbec for lipids analysis.

## Author contributions

Conceptualization: HM, JBM, DM, GM, JP.

Formal analysis: HM, LP, XG, DS, SN, ENH, JBM, DM, GM, JP.

Funding acquisition: HM, EM, JP.

Investigation: HM, LP, EM, XG, VV, PB, JR, AH, WS, JZ, GM.

Methodology: HM, VV, PB, DC, ENH, JBM, GM, JP.

Supervision: JP.

Visualization: HM, JP

Writing – Original draft: HM, JP.

Writing – Review and editing: All authors.

## Data availability

Global lipidomics data have been deposited in Zenodo (https://doi.org/10.5281/zenodo.10478862)

## STAR METHODS

### Key resources table

In a separate file

### Experimental models and subject details

#### Cells

*Aedes aegypti* Aag2, *Ae. albopictus* C6/36 (CRL-1660) and baby hamster kidney BHK-21 (CCL-10) cell lines from the American Type Culture Collection (ATCC) were grown in Roswell park memorial institute (RPMI) media (Gibco), supplemented with 10% heat-inactivated fetal bovine serum (FBS) (Gibco) and 1% Penicillin-Streptomycin (P/S) (Gibco) at 28°C for mosquito cells and at 37°C for BHK-21 with 5% CO_2_. Human hepatocellular carcinoma Huh-7 cell line (clone JTC-39) obtained from the Japanese Health Sciences Foundation, Osaka, Rhesus monkey kidney epithelial LLC-MK2 (CCL-7), and African green monkey kidney Vero (CCL-81) cell line from ATCC were maintained in Dulbecco’s modified Eagle medium (DMEM, Gibco), supplemented with 10% FBS, and 1% P/S at 37°C with 5% CO_2_. Mosquito cell media was supplemented with 1% non-essential amino acids (NEAA) (Thermo Fisher Scientific). Primary neonatal Human dermal fibroblasts (NHDF) (CC-2509, Lonza) were grown at 37°C with 5% CO_2_ in fibroblast growth basal medium (FBM, Lonza) supplemented with fibroblast growth medium-2 bulletkit (FGM-2, Lonza) and 2% FBS (Gibco). All cell lines were yearly tested negative for mycoplasma contamination with specific primers. Buffy coats from healthy donors (n=4) were obtained from the Etablissement Français du Sang (EFS, Montpellier, France). CD14+ monocyte isolation and differentiation to monocyte-derived dendritic cells (moDCs) were performed as previously described ^1,2^. MoDCs were maintained in RPMI supplemented with 10% FBS and 1% P/S at 37°C with 5% CO_2_.

#### Viruses

Dengue virus 2 (DENV 2) New Guinea C (NGC) strain and DENV2 16681 strain collected from a dengue fever patient in Thailand were obtained from the World Reference Center for Emerging Viruses and Arboviruses (WRCEVA) at UTMB, TX, USA. West Nile virus (WNV) strain IS98-ST1 ^3^ and chikungunya virus (CHIKV) LR2006_OPY1 strain isolated in la Réunion island were obtained from P. Desprès, University of la Réunion, France. Zika virus (ZIKV) PF-251013-18 strain was obtained via V. M. Cao-Lormeau and D. Musso, Institut Louis Malardé (ILM), Tahiti Island, French Polynesia. All viruses were propagated in C6/36 cells with 2% FBS. DENV productions were titrated using focus forming assay with BHK-21 cells, ZIKV and CHIKV productions were titrated using plaque assay with BHK-21 cells, and WNV was titrated using plaque assay with Vero cells. Viruses were stored at -70°C.

#### Mice

Four-week-old C57BL/6J male mice were purchased from Charles-River (France) and housed in ventilated cages in NexGen Mouse 500 (Allentown; Serial number: 1304A0078) in the biosafety level 3 (BSL-3) animal facility at MIVEGEC-IRD, Montpellier, France. Mice were maintained with 17h:7h light/dark cycle, 53-57% humidity, 20-24°C temperature and provided with irradiation-sterilized mouse diet (A03, SAFE, France) and sterilized water *ad libitum*. Mice were used for experiments one week after reception from Charles-River to let them accommodate to the animal facility. Every effort was made to minimize murine pain and stress. All animal protocols were approved by the APAFiS national ethical committee (permission number: 43466).

#### Mosquitoes

The BORA *Ae. aegypti* colony collected in Bora-Bora island in 1980 ^4^ was reared in the VectoPole insectary, MIVEGEC. Eggs were hatched in deionized water and larvae were fed grinded fish food (TertraMin, Tetra) at 26°C under 12h:12h light-dark cycle until pupation. Adult mosquitoes were kept in Bioquip cages at 28°C, 70% relative humidity with a 14h:10h light-dark cycle and access to 10% sugar water solution.

## METHOD DETAILS

### Isolation of extracellular vesicles (EVs)

EV purification was based on previously described protocols ^5^. 9 x 10^6^ Aag2 cells or 12 x 10^6^ Huh7 cells, covering 70-90% of T175 flask surface, were reared in complete media. Supernatant was replaced with FBS-free RPMI medium supplemented with 1% P/S and 1% NEAA (for mosquito cells), and collected after 48h. FBS-free, EV-containing supernatant was centrifugated at 300 x g for 10 min to remove large cell debris, at 2,000 x g for 10 min to remove small cell debris and at 10,000 x g for 30 min to remove large EVs. Supernatant fraction collected from the previous step was ultra-centrifugated at 100,000 x g for 155 min and the pellet was washed with ice-cold PBS (Gibco) before a second ultracentrifugation at 100,000 x g for 155 min to eliminate contaminants. The resulting pellet was resuspended in 800 µl of PBS for EV functional assay, in 20 µl of 0.2 % BSA PBS for the EV uptake assay or in 600-1500 µl 1X RadioImmunoPrecipitation Assay (RIPA) buffer (Thermo Scientific) for lipid extraction (see Table S2 for details). Proteins were quantified in EV lysates using Qubit Protein Assay kit (Invitrogen).

### Flow cytometric analysis of mosquito EV uptake by human cells

20 µl of the concentrated mosquito EVs or similarly obtained material from an equal volume of non-conditioned culture medium were stained with 1 µl of PKH26 (Sigma-Aldrich) in 200 µl of diluent C. EVs were purified through density gradient by mixing with 60% iodixanol (Optiprep; Axis-Shield) to a final concentration of 45% iodixanol and overlaid with a linear gradient of 40-5% iodixanol in PBS. Density gradients were centrifuged at 190,000 x g for 16 h in SW60 rotor. EV-containing gradient fractions (1.06 g/ml -1.08 g/ml) of 308 µl were collected and EVs were detected by high-resolution flow cytometry using a Cytek Aurora flow cytometer with Enhanced Small Particle module as previously detailed ^53,54^. The EV-containing gradient fractions were pooled, diluted with PBS + 0.1% EV-depleted BSA and centrifuged at 190,000 x g for 65 min at 4°C. The EV-containing pellet was resuspended in 150 µl of EV-depleted RPMI and 20 µl of this suspension was added to 10,000 Huh7 cells. At 2.5, 5 or 24h, cells kept at 4 or 37°C were detached via trypsinization with 0.05 % trypsin for 5 min at 37°C. Uptake of PKH26-labeled EVs by the Huh7 cells was assessed using a Cytek Aurora flow cytometer equipped with three lasers (Cytek Biosciences Inc) with conventional cell acquisition setting. Data analysis was performed using FlowJo v10.07 software (FlowJo LLC, Ashland, OR).

### Extraction of nucleotide-free EV-proteins and EV-lipids

Volumes of EV lysates containing approximately 110 µg of proteins were treated with 2.5 μl of DNAse I recombinant (10 Units/µl) (Roche Diagnostics), 5 μl of 10X DNase buffer, and 10 μl of RNAse A (10 µg/µl) (Thermo Scientific) followed by incubation for 45 min at 37°C. 1 mL of methanol:dichloromethane:nuclease-free water (2:1:1 volume) was then added. The resulting mixture was vortexed for 1 min and centrifuged at 700 x g for 6 min to pellet proteins. The pellet was resuspended in 50 µl of DMSO.

The monophasic liquid phase, containing lipids and other metabolites, was collected, dried under nitrogen and resuspended in 50 μl of DMSO (Sigma-Aldrich). 300 μl of methanol:chloroform (1:1 volume) was added before vortexing for 1h at l of 4°C. 50 μl nuclease-free water was added and the resulting solution was vortexed for 1 min and centrifuged at 1,800 x g for 10 min. The protein interphase was added to the previously resuspended protein pellet. The upper organic phase, containing lipids, was dried under nitrogen, resuspended in 50 μ of DMSO and stored at -70°C.

DNA, RNA and proteins in the different extracts were quantified using Qubit dsDNA High Sensitivity (HS), Qubit RNA Broad Range (BR) and Qubit Protein Assay kits (Invitrogen), respectively.

### Isolation of DENV for lipid extraction

Five T175 flasks of C6/36 or LLC-MK2 cells were incubated with DENV2 16681 strain at MOI of 1 for 1h at 28°C for C6/36 or 37°C for LLC-MK2 cells. Mock infections were similarly conducted. On day 3 for LLC-MK2 and day 6 for C6/36, total supernatant was collected. Virions and EVs were precipitated using 10% (w/v) PEG 8,000 (Merck) and 1.5 M NaCl (Merck) at 4°C overnight. After centrifugation at 12,000 x g, the pellet was resuspended in TNE buffer [10 mM Tris-HCl, pH 7.5, 140 mM NaCl, 1 mM EDTA (Bio Basic)] supplemented with 10% (w/v) sucrose (Merck) and layered onto a discontinuous sucrose step gradient, consisting of 30% and 60% sucrose (w/v), creating a 10/30/60 step gradient. Ultra-centrifugation was carried at 82,705 x g for 3.5 h and the fraction immediately above the 60% sucrose cushion was collected, diluted using TNE and pelleted via ultracentrifugation at 82,705 x g for 1.5 h. The resulting pellet was re-suspended in 100 µl of TNE buffer containing 1% BSA. Plaque assay on isolated fractions confirmed the recovery of viruses, as previously published ^6^. Purifications of DENV was performed in Prof. Duncan Smith’s Laboratory at Mahidol University, Bangkok, Thailand. Purified virions and vesicles were shipped in 100% methanol (Merck) on dry ice to IRD, Montpellier, France for lipid extractions as described above.

### Cell infection upon supplementation with EVs, EV-proteins or EV-lipids

In 24-well plates, 2.5 x 10^5^ Huh7, NHDF or moDC cells were incubated with DENV2 NGC, WNV, ZIKV or CHIKV at a MOI of 0.1 in 200 µl of FBS-free cell-corresponding media for 1h at 37°C. The inoculum was supplemented with either (i) 0.1 or 1 µl of concentrated EVs, (ii) different quantities of EV-proteins ranging from 0.051-4.85 µg, (iii) 0.01, 0.1 or 1 µl of EV-lipid extracts, (iv) 0.1 µl of EV-lipid fractions or (v) 0.01, 0.1 or 1 µl of a sphingomyelin (SM) solution, obtained by resuspending 25 mg of commercial SM pig brain powder in 5 ml of nuclease-free water. To homogenize volumes of DMSO used as vehicle for EV-protein extracts, EV-lipid extracts and EV-lipid fractions, the different supplemented volumes were complemented to 1 µl with DMSO. Controls for EVs and SM solutions were supplemented with 1 µl of nuclease-free water; controls for EV-lipid extracts were supplemented with 1 µl of DMSO; controls for EV-lipid fractions were supplemented with 0.1 µl of the corresponding blank extraction resuspended in 1 µl DMSO; and controls for EV-protein extracts were supplemented with 1 µl of DMSO. After incubation, 200 μl of 4% FBS cell-corresponding media was added. At 72h post-infection for flaviviruses and 48h for CHIKV, cells were lysed in 350 μl of TRK lysis buffer (EZNA RNA Kit I, Omega) and supernatant was collected.

### Focus Forming Unit (FFU) assay

100,000 BHK-21 cells per well of 24-well plates were incubated with 150 μl of 10-fold serial dilutions of inoculum for 1h at 37°C with l of sterile 2% CarboxylMethyl Cellulose (CMC) in RPMI supplemented with 2% FBS and 1% P/S was added. Three days later, cells were fixed with 4% paraformaldehyde (Sigma-Aldrich) for 30 min, permeabilized with 0.3% Triton X-100 (Sigma-Aldrich) PBS for 30 min, stained with pan-flavivirus 4G2 antibody (kindly provided by S. Vasudevan from Duke-NUS, Singapore) at 1:400 in 1% BSA (PAN-Biotech) for 1h at 37°C, and stained with secondary anti-mouse Alexa Fluor 488-conjugated antibody (Invitrogen) at 1:500 in 1% BSA for 1h at 37°C in the dark. Foci were counted using EVOS M5000 imaging system (ThermoFisher) and averaged over three replicates to calculate FFU/ml.

### Relative quantification of viral gRNA

Total RNA from cells was extracted using EZNA Total RNA kit I (Omega). Relative quantification of the positive strand of viral gRNA for DENV, ZIKV, WNV and CHIKV was obtained by two-step RT-qPCR. RNA extracts were treated with DNAse and reversed transcribed using gDNA Clear cDNA Synthesis kit (Bio-Rad). qPCR was conducted in 10 µl final volume with 2 μl of 5X HOT Pol EvaGreen qPCR mix plus (Euromedex), 300 nM of forward and μ reverse primers (Table S5) in AriaMx Real-time PCR System (Agilent) with the following thermal conditions: 95°C for 15 min, 45 cycles at 95°C for 15s, 60°C for 20s and 72°C for 20s, followed by a melting curve analysis. *GAPDH* mRNA levels were quantified by qPCR in the same conditions for normalization. Gene expression fold change was calculated by the ΔΔCq method.

### Evaluation of cell viability

At 72h post treatment, cell viability was estimated by calculating the inverse differences in GAPDH *Ct* relative to the control conditions and the number of viable cells was evaluated using CyQUANT NF Cell Proliferation Assay Kit (Invitrogen).

### Absolute quantification of DENV and WNV (+) gRNA

Total RNA was extracted from cell lysates using EZNA RNA extraction kit I or from blood samples using QIAamp Viral RNA Mini Kit (Qiagen). Absolute quantification of positive strand gRNA was performed through one-step RT-qPCR. Total reaction volume was 10 µl and contained 5 µl of iTaq Universal SYBR green one-step kit (Bio-Rad), 300 nM of forward and reverse primers (Table S5) and 2 μl of RNA extract. The reaction was performed in AriaMx Real-time PCR System with the following thermal profile: 50°C for 10 min, 95°C for 1 min and 40 cycles of 95°C for 10 sec and 60°C for 15 sec, followed by a melting curve analysis. An absolute standard curve for DENV and WNV gRNA was generated by amplifying the qPCR target using primers detailed in Table S5 as performed previously ^7,8^.

### Attachment and internalization assays

For attachment assay, 2.5 × 10^5^ Huh7 cells prechilled at 4 °C for 15 min were incubated with DENV2 NGC at a MOI of 0.1 in 200 μl of serum-free DMEM supplemented with 0.1 μl of EV-lipid extract complemented with DMSO to 1 µl or 1 µl of DMSO (control) for 30 min at 4 °C. Inoculum was removed and cells were washed thrice with prechilled 2 % FBS DMEM. Attached viruses were quantified as gRNA copies in cell lysates extracted using EZNA RNA extraction kit I. For internalization assay, Huh7 infection was repeated and non-internalized virus particles were removed by adding 2 mg/ml of pronase (Sigma-Aldrich) in 200 µl of serum-free medium for 5 min on ice. After two washes, internalized viruses were quantified as gRNA copies in cell lysate extracted with EZNA RNA extraction kit I.

### Translation assay

At 3, 4, 5, and 6h post-infection upon supplementation with 0.1 µl of EV-lipid extracts, Huh7 cells were washed twice with PBS, scrapped in 70 μl of RIPA with 1X protease inhibitors (Sigma-Aldrich) and supernatant was collected after centrifugation at 12,000 g for 1 min. Normalized protein quantities were separated under denaturing conditions in 10% polyacrylamide gel and transferred onto 0.2 µm Nitrocellulose membrane using TransBlot system (BioRad). Staining was conducted with 1:2,000 anti-DENV2 NS3 (GTX124252, Genetex) and 1:400 anti-Actin (MA5-11869, Invitrogen), and 1:2,000 of goat anti-rabbit (7074P2, Cell Signaling) and goat anti-mouse (7076P2, Cell Signaling) as secondary antibodies, respectively, in PBS-tween 0.1% with 1% BSA. Translation assay was repeated by adding 20 µM of NITD008 (Sigma-Aldrich) in cell media for 2h before infection and during the course of infection.

### Replication Assay

At 1, 3, 6, 9, 12, 15, 18, 21 and 24h post-infection, Huh-7 cells were washed with PBS and lysed in 350 μl of TRK lysis buffer. Total RNA was extracted using EZNA Total RNA extraction kit I and used for relative quantification of the negative strand of DENV gRNA [(-)gRNA] using two-step RT-qPCR with TaqMan probe. RNA extracts were treated with DNAse and reversed transcribed using gDNA Clear cDNA Synthesis kit (Bio-Rad). qPCR was conducted in total reaction volume of 10 µl containing 2 μl of cDNA, 5 μl of 2X iTaq Universal probe kit (Bio-Rad), 300 nM of forward and reverse primers and 200 nM of probe (Table S5) in AriaMx machine with the following thermal conditions: 95°C for 2 min followed by 45 cycles of 95°C for 10 sec and 60°C for 30 sec. *GAPDH* mRNA was quantified as above for normalization.

### Relative quantification of total nascent proteins

At 6h post-treatment (hpt), nascent proteins were quantified using Protein Synthesis Assay Kit (Abcam). Briefly, the media were replaced with 1X Protein Label solution and cells were incubated for 1.25 h at 37 °C. Negative control cells were neither exposed to the Protein Label solution not treatment, whereas positive control cells were incubated with 1X Protein Label solution without treatment. After wash with 100 μ Solution was added and incubated for 15 min at room temperature (RT) in the dark. After wash with 200 μl of 1X Wash Buffer (WB), cells were treated with 100 μl of 1X Permeabilization Buffer (PB) for 10 min at RT, then with 100 µl of 1X reaction cocktail (97 μl PBS, 1 μl of 100X Copper Reagent, 1 μl of 100X Fluorescent Azide and 1 μl 20X Reducing Agent) for 40 min at RT in the dark. Finally, after 2 washes in 200 μl of WB, 100 μl of PBS was added to each well before measuring fluorescence intensity at excitation/emission of 430(20)/535(35) nm using a Spark multimode microplate reader (TECAN). Relative quantities were obtained by subtracting values from negative controls and normalizing to cells treated with DMSO.

### Western blot quantification of IRE1α and phospho-IRE1α

At 6 hpt, cells were washed with PBS, scrapped in 70 µl of RIPA containing 1X protease and 1X phosphatase inhibitors. WB was conducted as described above for translation assay and staining was performed with 1:500 anti-IRE1α (#3294, Cell Signaling), 1:700 anti-pIRE1α (NB100-2323, Novus) and 1:400 anti-Actin (MA5-11869, Invitrogen), and 1:2,000 of goat anti-rabbit and goat anti-mouse as secondary antibodies.

### Relative quantification of ERAD and IFN-related genes

Total RNA from cells was extracted using EZNA Total RNA kit I. Expression for *Serl1 L, Derlin1, Edem1, Herpud1, Hrd1* and *GAPDH* was quantified through one-step RT-qPCR using iTaq Universal SYBR green one-step kit (Bio-Rad) with the corresponding primiers (Table S5) and the conditions described above for WNV (+)gRNA. Relative quantification was performed using the 2-ΔΔCt method by normalization to *GAPDH* Ct values.

Expression for IFN-related genes was conducted by reverse transcription using the PrimeScript RT Reagent Kit (Perfect RealTime, Takara Bio Inc.). qPCR reaction was performed in duplicate using Takyon ROX SYBR MasterMix blue dTTP (Eurogentec) on an Applied Biosystems QuantStudio 5 (Thermo Fisher Scientific) in 384-well plates. Transcripts were quantified using the following program: 3 min at 95°C followed by 35 cycles of 15 s at 95°C, 20 s at 60°C, and 20 s at 72°C. Values for each transcript were normalized to the geometric mean of Ct values of 4 different housekeeping genes (RPL13A, ACTB, B2M, and GAPDH), using the 2-ΔΔCt method. Primers used for quantification of transcripts by qPCR are listed in Table S5.

### Metabolite extraction from cells

Four hours post-treatment, Huh-7 cells were washed with 0.9% NaCl (Sigma) and 4 wells per condition were collected in 500 µl of ice-cold methanol (LC-MS grade, Thermo Fisher Scientific) and water in 80:20 ratio (v/v) by scraping, and sonicated in an ultrasonic bath (J.R. Selecta) for 15 min at 4 °C. Samples were centrifuged at 10,000 x g for 1 min at 4 °C, and 400 μl of supernatant were collected. Pellets were extracted a second time by adding 500 μl of methanol:water (80:20 volume) solution, sonicated and centrifuged before collecting another 400 μl of supernatant. Combined supernatants were dried, weighted and stored at -70 °C.

### Untargeted lipidomics

A Q Exactive Plus quadrupole (Orbitrap) mass spectrometer, equipped with a heated electrospray probe (HESI II) and coupled to a U-HPLC Vanquish H system (Thermo Fisher Scientific, Hemel Hempstead, U.K.) was used for Ultra-High-Performance Liquid Chromatography−High-Resolution Mass Spectrometry (UHPLC-HRMS) profiling. Dry extracts were normalized to 2 mg of dry mass/ml in an 80:20 (v:v) methanol:water solution. Separation was conducted using an Acquity UPLC CSH C18 (100 mm, 2.1 mm, 1.7 µm) equipped with a guard column (Waters SAS). The mobile phase A (MPA) consisted in a mixture of acetonitrile/water (60:40; v/v) with 10 mM ammonium formate and 0.1% formic acid. The mobile phase B (MPB) consisted in an acetonitrile/isopropanol (90:10; v/v) with 10 mM ammonium formate and 0.1% formic acid. The solvent gradient was set as follow: 40% to 43% MPB (0 - 2 min), 50% MPB (2.1 min) to 54% MPB (2.1 – 11.9 min), 70% MPB (11.9 - 12 min), 70% to 99% MPB (12 – 17.9 min), 99% MPB (17.9 – 19.9 min). The flow rate was set to 0.3 mL/min, the autosampler temperature was 5 °C, the column temperature was 55 °C, and injection volume was 1 μl. Mass detection was performed in positive ionization (PI) mode (MS1 resolution power = 35 000 [full width at half-maximum (fwhm) at 400 m/z]; MS2 resolution power = 17 500; MS1 automatic gain control (AGC) target for full scan = 1 x10^6^; 1x10^5^ for MS2). Ionization spray was set to a 3.5 kV voltage, and the capillary temperature was 256 °C. The mass scanning range was m/z 100−1500. Data-dependent acquisition of MS/MS spectra for the six most intense ions followed each full scan. Stepped normalized collision energy of 20, 40, and 60 eV was used for data acquisition in data dependent analysis mode.

MS-DIAL v. 4.80 ^9^ was used for UHPLC-HRMS raw data analysis. Mass feature extraction ranged between 100 and 1500 Da and 0.5 to 18.5 min. MS1 and MS2 tolerance in centroid mode were set to 0.01 and 0.05 Da, respectively. Optimized detection threshold was set to 10^6^ and 10 for MS1 and MS2, respectively. Peaks were aligned to a quality control (QC, aliquot of all sample extracts) reference file, with a retention time tolerance of 0.15 min and a mass tolerance of 0.015 Da. The LipidBlast internal MS-DIAL database was used for putative annotation. MS-CleanR workflow version 1.0 ^10^ was employed for cleaning MS-DIAL data. A minimum blank ratio of 0.8, a maximum relative standard deviation (RSD) of 40, and a relative mass defect (RMD) ranging from 50 to 3000 were set for all filters selected. For feature relationships detection, the maximum mass difference was set to 0.005 Da, and the maximum RT difference to 0.025 min. The Pearson correlation links were considered with correlation ≥ 0.8 and statistically significant with α = 0.05. The most intense and the most connected peaks were kept in each cluster. Feature not annotated within MS-DIAL were elucidated with MS-FINDER version 3.52 ^11^. The MS1 and MS2 tolerances were respectively set to 5 and 10 ppm. Formula finder was only processed C, H, O, N, P, and S atoms. The databases (DBs) were constituted from MS-FINDER internal DBs with LipidMaps and HMDB. Data were normalized by autoscaling before selecting regulated metabolites with more than two-fold intensity change and p-value < 0.1, as indicated by an unpaired t test with false-discovery rate (FDR) adjustment using MetaboAnalyst (v. 5.0) ^12^.

### Targeted quantitative lipidomics

Aag2 EVs isolated as detailed above were lysed in RIPA buffer. Quantification was performed for major phospholipids (PL) [including phosphatidylserine (PS), phosphatidylethanolamine (PE), phosphatidylcholine (PC) and phosphatidylinositol (PI)], major sphingolipids [including ceramide (Cer) and sphingomyelin (SM)], neutrals lipids (NL) [including cholesterol (Chol), cholesterol ester (Chol E) and triacylglycerol (TAG)] and total fatty acids (FA) [including saturated fatty acids (SAFA), monounsaturated fatty acid (MUFA) and polyunsaturated fatty acid (PUFA)]. 200 µg protein equivalent of EV lysates were extracted and analyzed differently for each lipid class.

For PL and NL, EV lysates were extracted according to Bligh and Dyer ^13^ in dichloromethane:water:methanol (2.5:2:2.5, v/v/v) with 2% acetic acid in the presence 100 µl of NL internal standards (stigmasterol; cholesteryl heptadecanoate; glyceryl trinonadecanoate) and 40 µl of PL internal standards (PC 13:0/13:0; Cer d18:1/12:0; PE 12:0/12:0; SM d18:1/12:0; PI 17:0/14:1; PS 12:0/12:0). Samples were centrifugated at 500 x g for 6 min, evaporated to dryness and resuspended in 20 µl of ethyl acetate for NL analysis and 50 µl of methanol for PL analysis. For NL analysis, 1 µl of extract was analyzed by gaz-chromatography flame-ionisation-detector (GC-FID) on a GC TRACE 1300 Thermo Electron system using an Zebron ZB-5MS Phenomenex columns (5% polysilarylene, 95% polydimethylsiloxane, 5m X 0.25 mm i.d, 0.25 µm film thickness) ^14^. Oven temperature was programmed from 190°C to 350°C at a rate of 5°C/min and the carrier gas was hydrogen (5 ml/min). The injector and the detector were at 315°C and 345°C, respectively. For PL analysis, 2 µl of extract was analyzed using an Agilent 1290 UPLC system coupled to a G6460 triple quadripole mass spectrometer (Agilent Technologies). A Kinetex HILIC column (Phenomenex, 50 x 4.6 mm, 2.6 µm) was used for LC separations. The column temperature was controlled at 40°C. The mobile phase A was acetonitrile and B was 10 mM ammonium formate in water at pH 3.2. The gradient was as follow: from 10% to 30% B in 10 min; 10-12 min, 100% B; and then back to 10% B at 13 min for 2 min prior to the next injection. The flow rate of mobile phase was 0.3 ml/min. Electrospray ionization was performed in positive mode for Cer, PE, PC and SM analysis and in negative mode for PI and PS analysis. Needle voltage was set respectively at 4 kV and -3.5 kV. Approximate quantification was obtained for each species through comparison to the internal standards of the concerned lipid family.

For total FA, approximate quantification concerns conventional FA: c10:0, c12:0, c14:0, c15:0, c16:0, c17:0, c18:0, c20:0, c22:0, c23:0, c24:0, c14:1w5, c15:1, c16:1w7, c18:1w9, c18:1w7, c20:1w9, c22:1w9, c24:1w9, c18:2w6, c18:3w6, c18:3w3, c20:2w6, c20:3w3, c20:3w6, c20:4w6, c20:5w3, c22:2w6, c22:6w3, c22:4w6. EV lysates were extracted as described above according to Bligh and Dyer method in the presence of internal controls (TAG19). Samples were centrifugated at 500 x g for 6 min, hydrolyzed in KOH (0.5 M in methanol) at 55°C for 30 min, and transmethylated in 14% boron trifluoride methanol (Sigma) and heptane (Sigma) at 80°C for 1h. Water and heptane were added. Samples were centrifugated at 500 x g for 1 min, dried and resuspended in 20 µl of ethyl acetate. 1 µl of extract was analyzed by GC-FID ^16^ on a Clarus 600 Perkin Elmer system using a Famewax RESTEK fused silica capillary columns (30 m x 0.32 mm i.d, 0.25 µm film thickness). Oven temperature was programmed from 100°C to 250°C at a rate of 6°C/min and the carrier gas was hydrogen (1.5 ml/min). The injector and the detector were at 220°C and 230°C respectively.

Peak detection, integration and quantitative analysis were done using Mass Hunter Quantitative analysis software (Agilent Technologies) based on quantity of internal standard.

### Lipid fractionation

NH2 cartridge HyperSep 500 mg (Thermo Fisher Scientific) were conditioned by adding 2 ml of chloroform:methanol (23:1 volume) followed by 2 ml of diethyl ether. 2 x 100 µl of EV-lipid extracts resuspended in diethyl ether were loaded into the cartridge sequentially. Solvents with increasing polarity were used to elute different classes of lipids from the cartridge, as previously described ^17^. Briefly, the 6 solvents used were: 2 ml of diethyl ether (F1), 1.6 ml of chloroform:methanol (23:1 volume) (F2), 1.8 ml of diisopropyl ether:acetic acid (98:4 volume) (F3), 2 ml of acetone:methanol (9:1.2 volume) (F4), 2 ml of chloroform:methanol (2:1 volume) (F5) and 2 ml of methanol with 0.2 M of ammonium acetate (F6). Blanks were generated by eluting an empty cartridge with the same solvents. Each fraction was dried before resuspension in 50 µl of DMSO.

### Sphingomyelin quantification in EVs from mosquito cells and mosquito saliva

Mosquito and human EVs were isolated as described above by ultracentrifugation and resuspended in 50 and 60 µl of SM assay buffer, respectively. Two pools of saliva were collected by letting 400 and 825 female mosquitoes feed on a Hemotek feeding system (Discovery Workshops) containing 3 ml of PBS. The saliva solutions were ultracentrifugated at 100,000 x g for 155 min and the pellets were resuspended in 30 and 60 µl of SM assay buffer, respectively. Absolute SM quantification was obtained with Sphingomyelin Quantification Colorimetric Assay Kit (Abcam) using an absolute standard equation generated from the kit reagents. 10 µl of cell EV solution (containing 74 and 62.4 µg of proteins and originating from an estimated cell number of 9 x 10^5^ and 4 x 10^6^ for Aag2 and Huh7 cells, respectively) or 7 and 10 µl of salivary EV solution (corresponding to salivas from 93 and 137.5 mosquitoes) from each saliva pools were quantified with the SM assay. To exceed the kit detection threshold, 1 nmol of standard SM was added to each sample. After adjusting sample volume to 50 µl with SM assay buffer, 34 µl SM Assay Buffer, 2 µl Sphingomyelinase, 10 µl ALP Enzyme, 2 µl SM Enzyme Mix, and 2 µl OxiRed Probe were added. Plates were incubated at 37°C for 2h before measuring absorbance at 570 nm using a Spark multimode microplate reader (TECAN). SM quantities in the samples were calculated by subtracting values for 1 nmol of SM standard.

### Sphingomyelinase (SMase) treatments

1µl of commercial SM solution (5 µg/µl), of mosquito EV-lipid extract or of EV-lipid fraction 6 was treated with 1 U of SMase (Sigma) for 1h at 37°C, then heated for 30 min at 65°C to inactivate SMase. As control, we similarly treated cell media with SMase. The resulting solution was used for supplementation during infection as described above.

### WNV injection in mice

Mice were shaved with 0.4 mm animal trimmer (VITIVA MINI, BIOSEB) on the lower back one day prior injection to limit any effect of shaving-induced inflammation. Mice anesthetized by injection of 0.2 ml/mouse solution, containing 10 mg/ml of ketamine (Imalgène 1000, Boehringer Ingelheim Animal Health) and 1 mg/ml of xylazine (Rompon 2%, Elanco GmbH), were intradermally (ID) inoculated with 10^3^ PFU of WNV mixed with 0.1 or 1 µl of mosquito EV-lipid extract complemented to 1 µl with DMSO. Total volume injected was complemented to 10 µl with PBS. Control mice were injected with 1 µl of DMSO with or without WNV inoculum in a total volume of 10 µl complemented with PBS. Mice were weighed before injection and daily thereafter to calculate the percent of weight loss. Daily clinical examinations were conducted and a clinical score (CS) ranging from 0 to 5 was assigned to each mouse following criteria from a previous study ^18^, where CS of 0 has been assigned to healthy mice; CS of 1 for mice with ruffled fur, lethargy, hunched posture, no paresis, normal gait; CS of 2 for mice with altered gait, limited movement in 1 hind limb; CS of 3 for lack of movement, paralysis in 1 or both hind limbs, and CS of 4 for moribund mice. A CS of 5 indicated mortality. At days 2, 4, 6, 8, and 10, blood samples were collected via mandibular puncture and sample volumes were estimated by pipetting. Mice were euthanized if they displayed neurological symptoms, severe distress, or weight loss exceeding 20%. Mice were euthanized under anaesthesia at day 12.

### Statistical analysis

Differences in relative gRNA copies were tested with one-tailed t-tests on log-transformed data to meet normal distribution. Differences in SM concentration gene expression were tested with one-tailed t-test or multiple LSD tests. A two-way repeated-measures ANOVA was used to test differences in weight changes, clinical score and RNAemia in mice. Differences in survival were tested with Kaplan-Meier survival analysis with Log-rank (Mante-Cox) comparison test. These statistical analyses were performed using Prism 8.0.2 (GraphPad).

