## Supplementary material for "Sphingomyelins in mosquito saliva modify the host lipidome to enhance transmission of flaviviruses by promoting viral protein levels": Key Ressources

**Key ressource table**

| Reagent or Resource | Source | Identifier, reference or Catalog number |
| --- | --- | --- |
| Experimental models : Cell lines | | |
| *Aedes aegypti* Aag2 | Provided by D. Missé, MIVEGEC, IRD, Montpellier |  |
| *Aedes albopictus* C6/36 | American Type Culture Collection [ATCC] | CRL-1660 |
| Human hepatocellular carcinoma Huh7 | Japanese Health Sciences Foundation, Osaka | JTC-39 |
| Baby Hamster Kidney BHK-21 | American Type Culture Collection [ATCC] | CCL-10 |
| African green monkey kidney Vero | American Type Culture Collection [ATCC] | CCL-81 |
| Neonatal Human Dermal Fibroblasts NHDF | Lonza | CC-2509 |
| Monocyte-derived dendritic cells (moDCs) | Produced by S. Nisole Lab, IRIM, CNRS, Montpellier |  |
| Rhesus monkey kidney epithelial LLC-MK2 | American Type Culture Collection [ATCC] | CCL-7 |
| Experimental models : Organisms/strains | | |
| Male mice : C57BL/6J | Charles River Laboratories (France) | Cat #C57BL/6J |
| Irradiation-sterilized mouse diet | A03, SAFE, France |  |
| Animal trimmer | VITIVA MINI, WAHL, BIOSEB, Germany | 1584-1693 |
| Imalgène 1000 | Boehinger Ingelheim Animal Healt, France | Code: VETO109AA301U |
| Rompon 2% | Elanco GmbH, Germany | Code : VETO109AA309U |
| Virus strains | | |
| Dengue virus 2 (DENV2)  New Guinea C (NGC) | World Reference Center for Emerging Viruses and Arboviruses (WRCREVA) at the University of Texas Medical Branch (UTMB) |  |
| DENV2 strain 16681 | D. Smith Lab, Mahidol University, Bangkok, Thailand  (Halstead et Simasthien 1970) |  |
| WNV strain IS98-ST1 | V. M. Cao-Lormeau and D. Musso, Institut Louis Malardé [ILM], Tahiti Island, French Polynesia |  |
| ZIKV PF-25013-18 | V. M. Cao-Lormeau and D. Musso, Institut Louis Malardé [ILM], Tahiti Island, French Polynesia |  |
| CHIKV | P. Duprès, Université de la La Réunion, France |  |
| Oligonucleotides and PCR-based reagents | | |
| Primers for qPCR amplification | Eurofins | Table S5 |
| iTaq Universal SYBR Green One-Step Kit | Biorad | 1725151 |
| iScript™ gDNA Clear cDNA Synthesis Kit | Biorad | 1725035 |
| iTaq Universal Probes One-Step Kit | Biorad | 1725141 |
| EvaGreen qPCR Mix | Euromedex | 08-25-00001 |
| PrimeScript RT Reagent Kit | Perfect RealTime, Takara Bio Inc. | RR037B |
| Takyon ROX SYBR MasterMix blue dTTP | Eurogentec | A58667 |
| Antibodies | | |
| Mouse Anti-enveloppe (4G2) | Provided by S. Vasudevan from Duke-NUS, Singapore |  |
| Goat Anti-mouse IgG (Alexa Fluor 488) | Life Technologies | A11029 |
| Rabbit anti-DENV2 NS3 | Genetex | GTX124252 |
| Mouse anti-Actin | Invitrogen | MA5-11869 |
| Anti-rabbit IgG, HRP-linked | Cell Signaling | 7074P2 |
| Anti-mouse IgG, HRP-linked | Cell Signaling | 7076P2 |
| Rabbit Anti-IRE1α | Cell Signaling | #3294 |
| Rabbit Anti-Phospho-IRE1α | Novus | NB100-2323 |
| Chemicals and enzymes | | |
| DMEM, high glucose, GlutaMAX™ Supplement, pyruvate | Gibco | 31966021 |
| RPMI 1640 Medium, GlutaMAX™ Supplement | Gibco | 61870010 |
| DPBS (1X) | Thermo Fisher Scientific | 14190250 |
| FBS | Eurobio | CVFSVF00 01 |
| MEM Non-essential amino acids (NEAA) | Thermo Fisher Scientific | 11350912 |
| Penicillin-Streptomycin (P/S) | Thermo Fisher Scientific | 11548876 |
| Trypsin - EDTA 0,25% | Thermo Fisher Scientific | 25200056 |
| Fibroblast Growth Medium-2 bulletkit (FGM-2) | Lonza | Cat # CC-3132 |
| ReagentPack^TM^ Subculture Reagents | Lonza | Cat # CC-5034 |
| Protease inhibitor cocktail | Roche | 11836170001 |
| Phosphatase inhibitor cocktail II | ThermoScientific | J61022 |
| 4% paraformaldehyde | Sigma-Aldrich | 47608-250ML-F |
| Methanol | Thermo Fisher Scientific | 10010280 |
| Ethanol | Thermo Fisher Scientific | 10680993 |
| Chloroform | Sigma-Aldrich | C2432-500ML |
| Diethylether | J.B-Michel Lab, MetaboHUB-MetaToul, I2MC, INSERM, Toulouse |  |
| Di-isopropylether | J.B-Michel Lab, MetaboHUB-MetaToul, I2MC, INSERM, Toulouse |  |
| DMSO | Sigma-Aldrich | D4540-100ML |
| Pronase | Sigma-Aldrich | 11459643001 |
| Sphingomyelinase (SMase) | Sigma-Aldrich | S8633 |
| RNase A, DNase and protease-free | Thermo Fisher Scientific | EN0531 |
| DNase I recombinant, RNase free | Roche | 04716728001 |
| NITD008 | Sigma-Aldrich | SML2409-5MG |
| Tween 20 | Sigma-Aldrich | P1379-500ml |
| MES SDS NuPAGE | Thermo Fisher Scientific | NP0002-02 |
| PageRuler™ Plus Prestained Protein Ladder | Thermo Fisher Scientific | 26620 |
| RIPA | Thermo Fisher Scientific | Cat # 89901 |
| Methyl Cellulose | Sigma-Aldrich | M0512-250G |
| BSA | PAN-Biotech | P06-1391100 |
| 40% Acryl/Bisacryl | Euromedex | EU0061-B |
| Trizma® Base | Sigma-Aldrich | T1503-500G |
| SDS 20% | Euromedex | EU0660-B |
| APS | Euromedex | EU0009-B |
| TEMED | Sigma-Aldrich | T9281-100ML |
| Trans-Blot Turbo RTA Mini 0.2 µm Nitrocellulose  Transfer Kit | Biorad | 1704270 |
| SuperSignal™ West Pico PLUS Chemiluminescent Substrate | Thermo Fisher Scientific | 34580 |
| Triton X-100 | Sigma-Aldrich | X100-100ML |
| Methanol (HPLC grade) | Fisher Chemicals, Waltham, MA USA | M/4056/15 |
| Acetonitrile (LCMS grade) | Fisher Chemicals Waltham, MA USA | A/0638/15 |
| Ammonium formate | Hipersolc Chromasolv (VWR chemicals) | 84884.180 |
| Formic acid (LCMS grade) | Hipersolc Chromasolv (VWR chemicals) | 84865.260 |
| Isopropanol | Fisher Chemicals Waltham, MA USA | P/7500/15 |
| Dichloromethane | 32222-2,5L / Fisher | Dichloromethane |
| Methanol | 15631400/ Fisher | 2LT Methanol LCMS |
| Acetic acid | 33209-1L/ Merck-Sigma | Acetic acid |
| NL internal standards (stigmasterol; cholesteryl heptadecanoate; glyceryl trinonadecanoate) | Stigmasterol : 1121-k MATREYA  Cholesterol ester C17: BJ492A Interchim  TG57: [T4632](https://www.sigmaaldrich.com/FR/fr/product/sigma/t4632) Merck | cholesteryl  heptadecanoate  glyceryl trinonadecanoate |
| PL internal standards (PC 13:0/13:0; Cer d18:1/12:0; PE 12:0/12:0; SM d18:1/12:0; PI 17:0/14:1; PS 12:0/12:0) | PC 13:0/13:0: 850340P / Merck  Cer d18:1/12:0:860512P / Merck  PE 12:0/12:0: 850702P / Merck  SM d18:1/12:0: 860583P / Merck  PI 17:0/14:1: 791641C / Merck  PS 12:0/12:0: 840038P / Merck |  |
| Ethyl acetate | 34972-2,5L/Fisher | Ethyl acetate |
| Ammonium formate | 73594-25G-F/ Merck | AMMONIUM ACETATE |
| deuterium-labeled internal standard [LxA4-d5, LTB4-d4, 5-HETE-d8 (Cayman Chemicals)] | LxA4-d5:10007737/ CAYMAN  LTB4-d4: 320110/CAYMAN  5-HETE-d8: 334230/ CAYMAN | 5(S),6(R)-LIPOXINA4-D5 Article AYR811 1X25ug  LEUKOTRIENE B4-d4  5(S)-Hydroeicosatetraenoic acid- d8 |
| Internal controls (TG17 or TG19 or TG15) | TG 19 T4632/Sigma | Glyceryl trinonadecanoate |
| Boron trifluoride methanol | 15716-1L /Merck | BORON TRIFLUORIDE METHANOL SOLUTION |
| Heptane | 15624770/ Fisher | 2.5LT Heptane LC-MS CHROMASOLV |
| Acetonitrile | A955­212/ Fisher | 2.5LT Acétonitrile, Optima |
| OptiPrep™ | STEMCELL Technologies | 17109821 |
| PKH26 | SIGMA-ALDRICH | MINI26 |
| Systems and Others | | |
| NH2 cartridge HyperSep™ 500mg | Thermo Fisher Scientific | 60108-518 |
| OASIS HLB 96-well plate (30 mg/well, Waters) | WAT058951/ UGAP | OASIS HLB 96-well Plate 30 mg Sorbent per Well 30 µm taille particule Reversed-Phase 0 - 14 |
| NexGen™ Mouse 500 | Allentown | 1304A0078 |
| EVOSTM M5000 microscope | Thermo Fisher Scientific | AMF5000 |
| Applied Biosystems QuantStudio 5 | Thermo Fisher Scientific | A34322 |
| AriaMx Real-time PCR System | Agilent | G8830A |
| Oa-Sys Heating System N-Evap 112 Nitrogen Evaporator | Organomation Associates |  |
| Q Exactive Plus quadrupole (Orbitrap) mass spectrometer | Thermo Fisher Scientific, Hemel Hempstead, U.K. | SN03123L |
| electrospray probe (HESI II) | Thermo Fisher Scientific, Hemel Hempstead, U.K. | 0924783A |
| U-HPLC Vanquish H system | Thermo Fisher Scientific, Hemel Hempstead, U.K. | 8324985 |
| Clarus 600 Perkin Elmer system with FID |  |  |
| Famewax RESTEK fused silica capillary columns (30 m x 0.32 mm i.d, 0.25 µm film thickness) | 12498/ Restek | Colonne capillaire Famewax, L 30m, DI 0.32mm, EF 0.25µm. |
| Acquity UPLC CSH C18 1.7 µm | Waters SAS, Guyancourt, France | 0178321671 |
| Guard column (CSH C18) | Waters SAS, Guyancourt, France | 0177321311 |
| ZorBAX SB-C18 column (2.1 mm, 100 mm, 1.8 µm) | RRHD SB-C18, 2,1x100mm, 1,8um, 1200 bars/ Agilent |  |
| GC TRACE 1300 Thermo Electron system with FID | N° de série 717001575/ Thermo Electron | Trace 1300- FID |
| RTX-5 | 10223/ Restek | Colonne capillaire Rtx-5, L 30m, DI 0.25mm, EF 0.25µm |
| Agilent 1290 UPLC system coupled to a G6460 triple quadripole mass spectrometer |  |  |
| Kinetex HILIC column ([Kinetex 2.6 µm HILIC 100 Å, LC Column 50 x 4.6 mm, Ea](https://www.phenomenex.com/products/kinetex-hplc-column/kinetex-hilic)) | 00B-4461-E0/ Phenomenex |  |
| Critical commercial assays | | |
| Qubit^TM^ dsDNA High Sensitivity kit | Thermo Fisher Scientific | Q32851 |
| Qubit^TM^ RNA Broad Range (BR) kit | Thermo Fisher Scientific | Q10210 |
| Qubit^TM^ Protein kit | Thermo Fisher Scientific | Q33211 |
| QIAamp Viral RNA Mini Kit | Qiagen | 52906 |
| EZNA Total RNA kit I | OMEGA | R6834-03 |
| Sphingomyelin Quantification Colorimetric Assay Kit | Abcam | ab287856 |
| CyQUANT NF Cell Proliferation Assay Kit | Invitrogen | C35007 |
| Protein Synthesis Assay Kit | Abcam | ab239725 |
| Sphingomyelin Assay Kit | Abcam | ab133118 |
| Software | | |
| GraphPad Prism |  | V8.0.2 |
| MS-DIAL | (Tsugawa et al. 2015) | Version 4.80 |
| MS-CleanR | (Fraisier-Vannier et al. 2020) | Version 1.0 |
| MS-FINDER | (Tsugawa et al. 2016) | Version 3.52 |
| MetaboAnalyst | (Pang et al. 2021) | Version 5.0 |
| Image Lab | Biorad | Version 6.1 |
| Aria Real-Time PCR | Agilent | Version 2.0 |
| Plastics | | |
| 24-well plates | Thermo Fisher Scientific | 142475 |
| 28-well plates | Thermo Fisher Scientific | 150687 |
| Cell culture flask, T-175 | SARSTEDT | 833912002 |
| Cell culture flask, T-75 | SARSTEDT | 833911002 |
| 0.2 ml non-skirted low profile 96 well PCR plate | Thermo Fisher Scientific | AB-0700 |
| Empty Fastprep® tubes | MP Biomedicals | 5076400 |
| Glass beads 1mm | Biospec Products | 11079110 |
