## Supplemental information for "Sphingomyelins in mosquito saliva modify the host lipidome to enhance transmission of flaviviruses by promoting viral protein levels"

Table of content

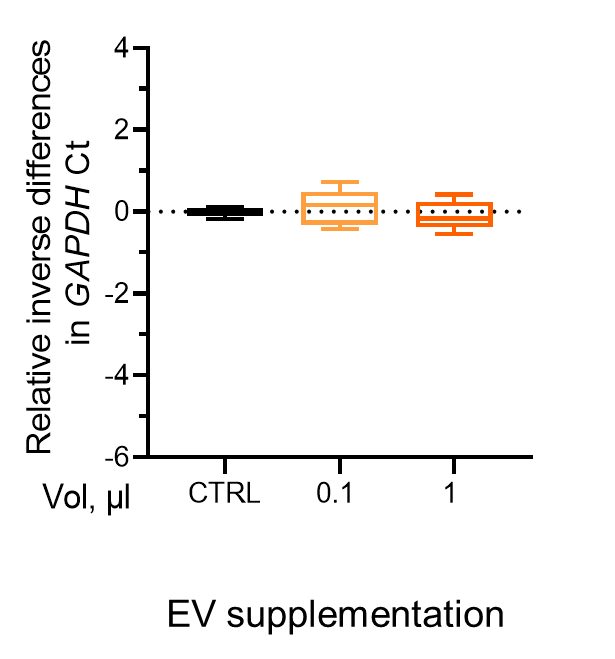

### Figure S1. Mosquito EVs do not alter cell viability

Huh7 cells were infected with DENV at MOI 0.1 upon supplementation with 0.1 and 1 µl of mosquito EVs. Cell viability was evaluated by calculating relative inverse differences in *GAPDH* Ct values. Results are presented as Tukey box and whiskers derived from at least 6 biological replicates collected from multiple experiments. CTRL indicates supplementation with 1X PBS.

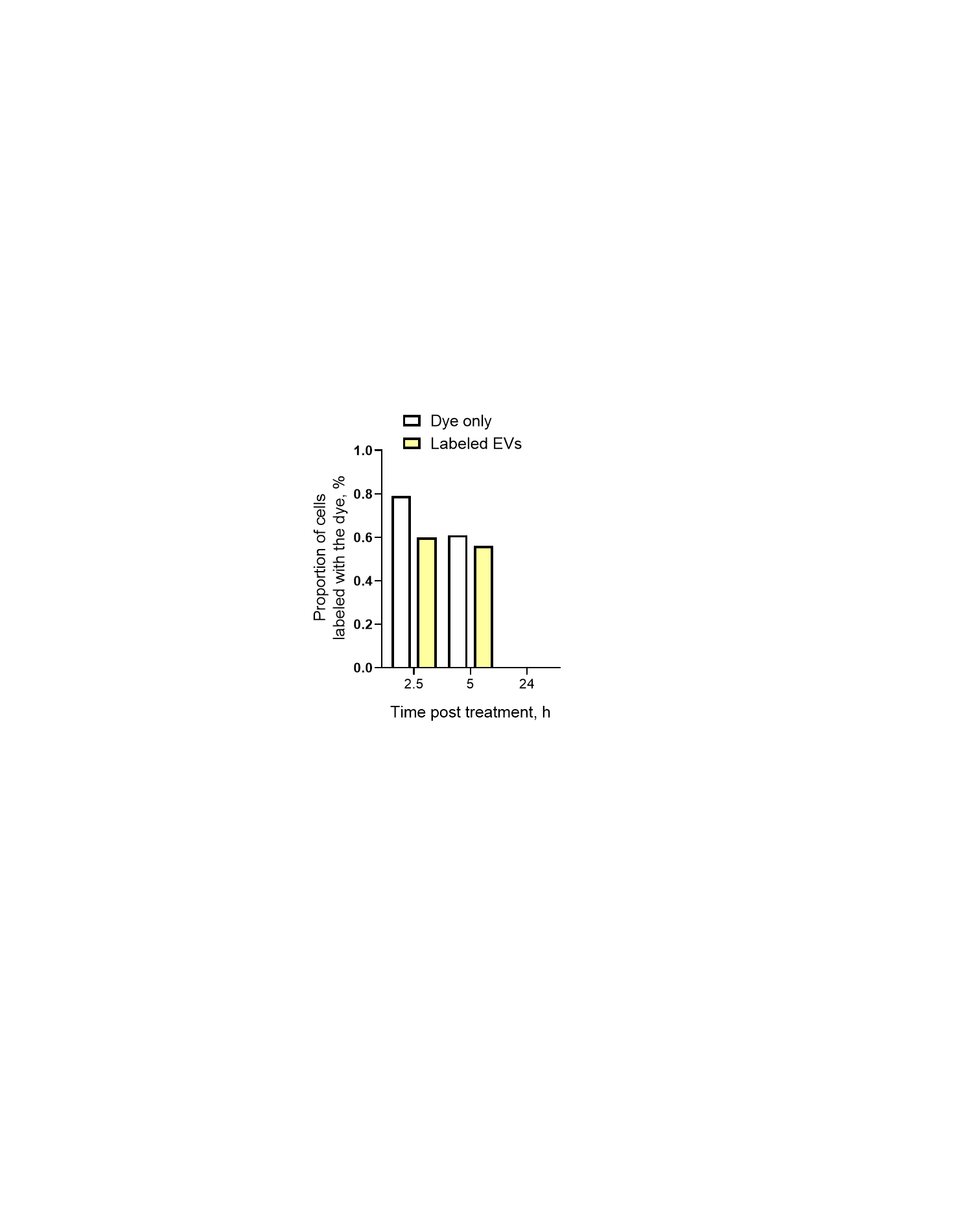

### Figure S2. Internalization of labelled EVs at 4°C

Huh7 cells were exposed to mosquito EVs labelled with PKH26 lipid dye at 4°C and dye internalization was quantified 2.5, 5 and 24h later. As controls, cells were exposed to dye-labeled material from an equal volume of non-conditioned medium.

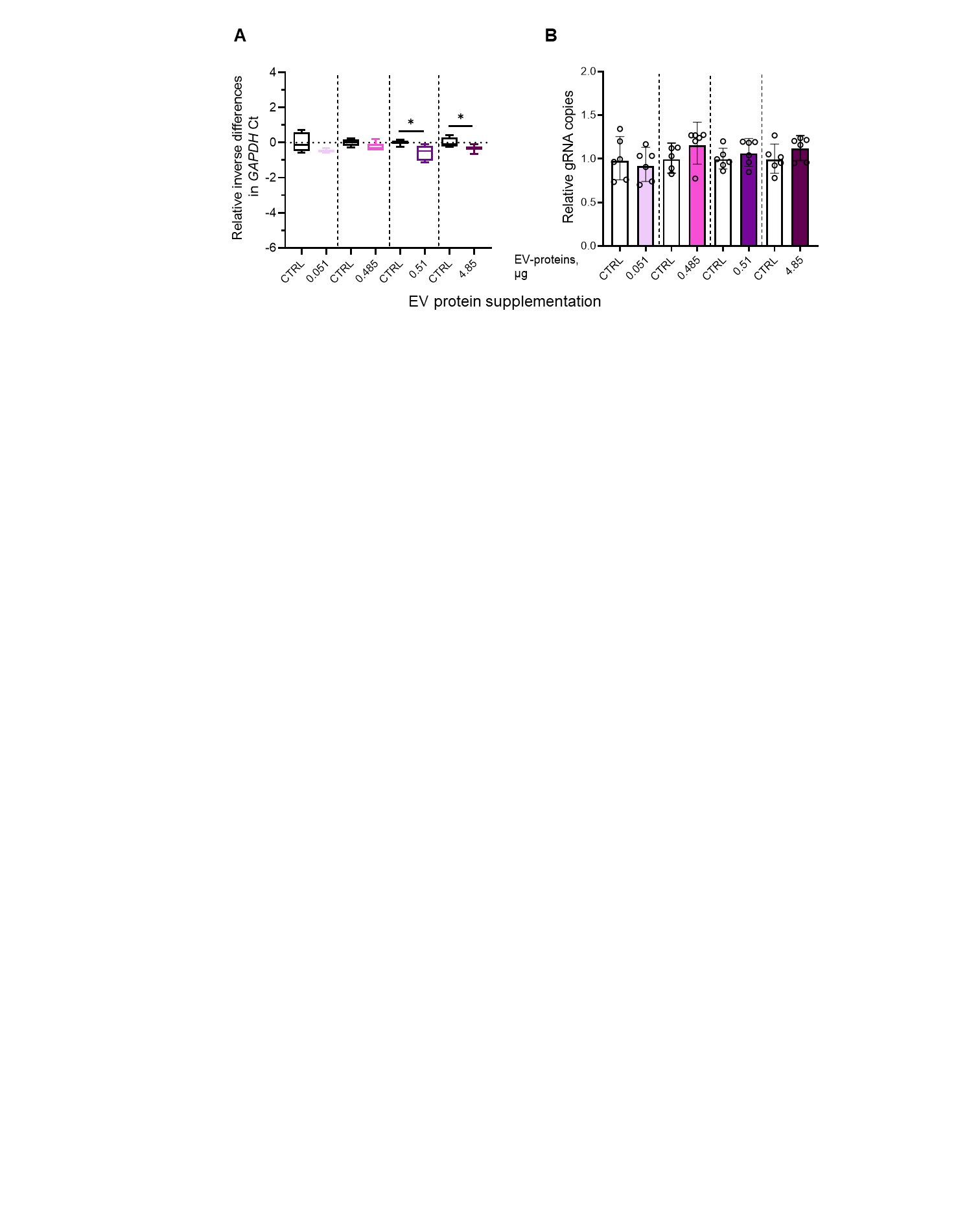

### Figure S3. Mosquito EV-proteins have no impact on DENV infection

Huh7 cells were infected with DENV at MOI 0.1 upon supplementation with 0.051, 0.485, 0.51, and 4.85 µg of protein extracts from mosquito EVs. Controls (CTRL) were supplemented with equivalent volumes of DMSO. ID of EV-protein extracts that were used: Prot 6.1 and Prot 6.2 as in Table S1.

(A) Cell viability was evaluated by calculating relative inverse differences in *GAPDH* Ct values. Results are presented as Tukey box and whiskers.

(B) Intracellular gRNA was quantified at 72 hpi for DENV. Bars show geometric means ± 95% C.I..

(A, B) Results derived from at least 6 biological replicates collected from multiple experiments. Points represent biological repeats. *, p < 0.05.

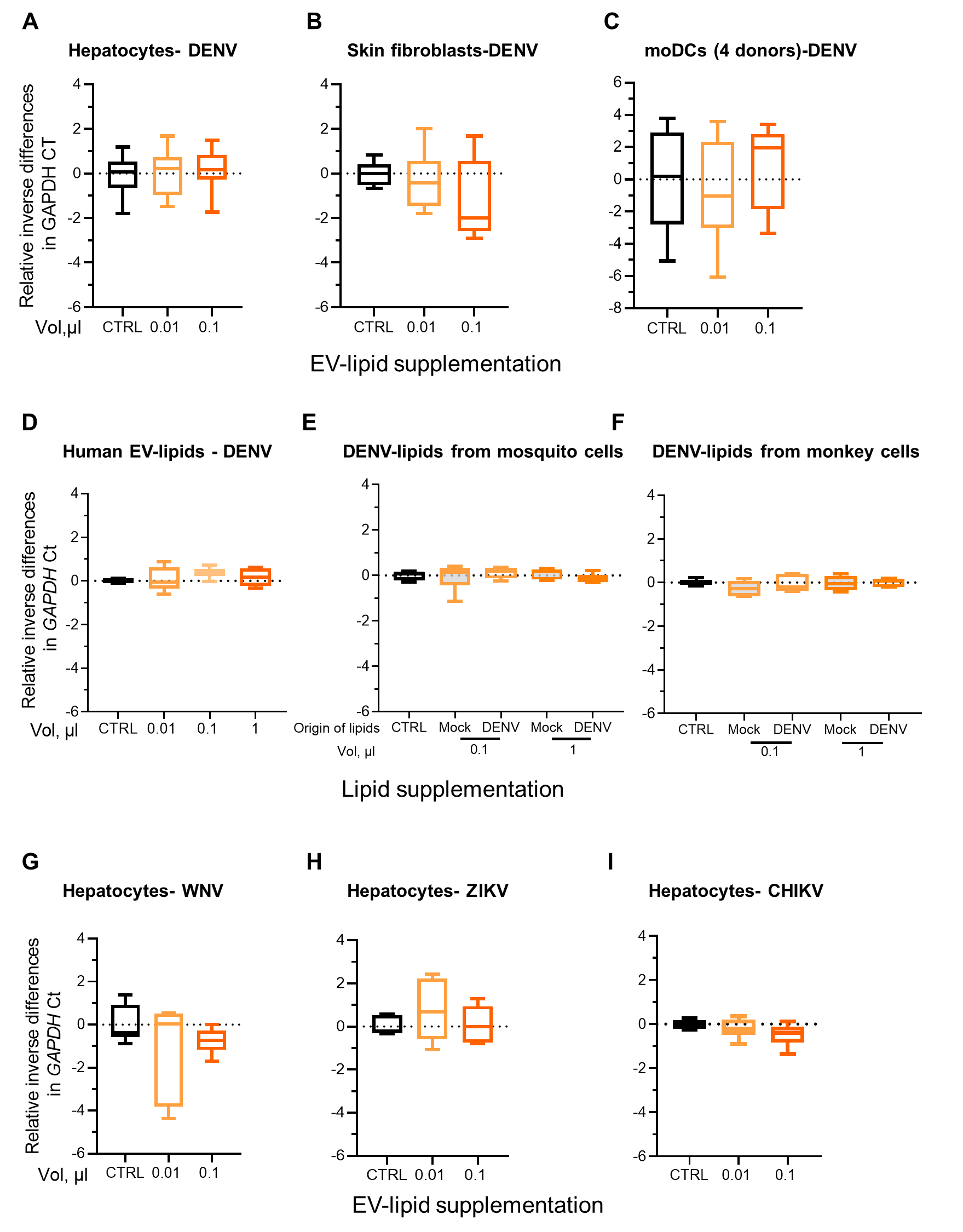

### Figure S4. Mosquito EV-lipids do not alter cell viability

(A-C) Human hepatocytes (Huh7) (A), primary neonatal human skin fibroblasts (NHDF) (B) and monocyte derived dendritic cells (moDC) (C) were infected with DENV at MOI 0.1 upon supplementation with 0.01 or 0.1 µl of mosquito EV-lipid extract.

(D) Huh7 cells were infected with DENV at MOI 0.1 upon supplementation with 0.01, 0.1 and 1 µl of human EV-lipids.

(E, F) Huh7 cells were infected with DENV at MOI 0.1 upon supplementation with 0.1 or 1 µl of DENV lipid extracts produced from mosquito (E) or mammalian (F) cells. Mock indicates supplementation with lipid extracts from the same DENV density fraction from mosquito (E) and mammalian (F) mock-infected cells.

(G-I) Huh7 cells were infected with WNV (G), ZIKV (H) or CHIKV (I) at MOI of 0.1 upon supplementation with 0.01 or 0.1 µl of mosquito EV-lipids.

(A-I) Cell viability was evaluated by calculating relative inverse differences in *GAPDH* Ct values. Results are presented as Tukey box and whiskers derived from at least 6 biological replicates collected from multiple experiments. CTRL indicates supplementation with DMSO.

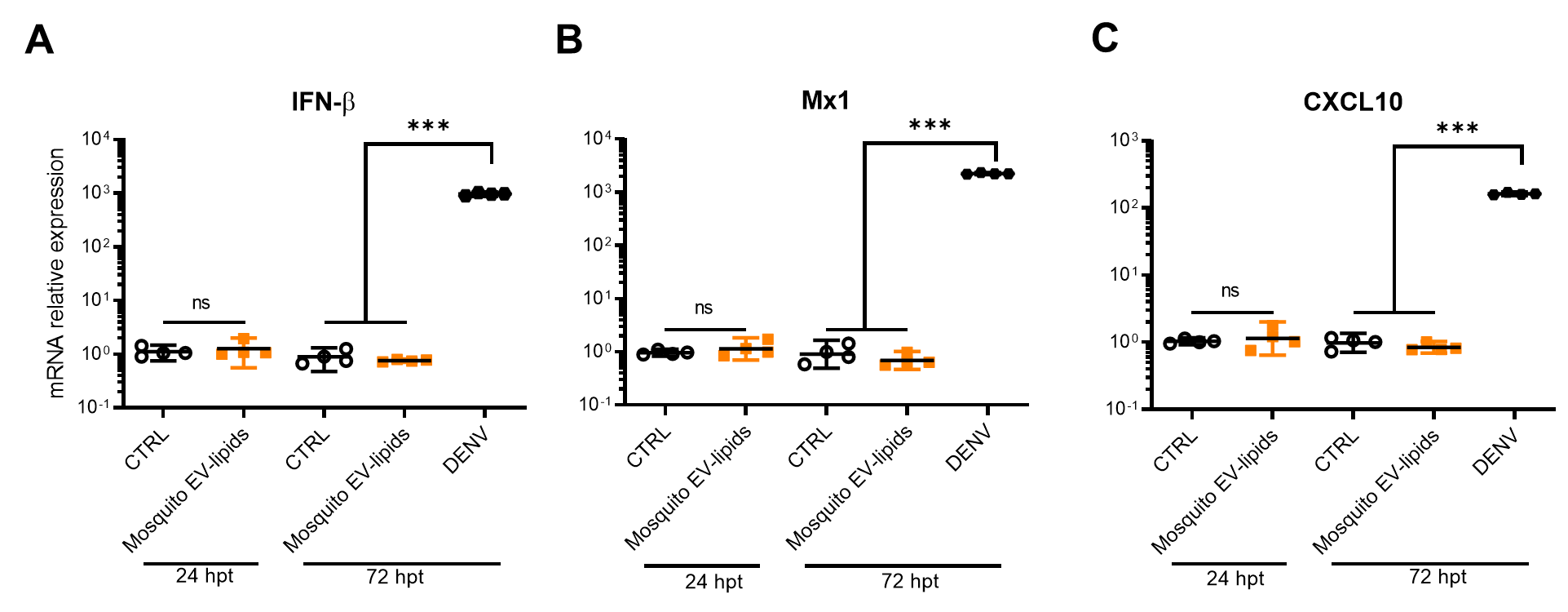

### Figure S5. Mosquito EV-lipids do not induce innate immune response

(A-C) Huh7 cells were supplemented with 0.1 µl of mosquito EV-lipid extracts or infected with DENV at MOI 0.1. *IFN-β* (A), *MX1* (B), and *CXCL10* (C) relative expressions were quantified at 24 and 72 hours post treatment (hpt). Bars show means ± SEM. Points represent biological repeats. CTRL indicates supplementation with DMSO. Ns, non-significant; ***, p-value < 0.001, as determined by one-tailed t-test.

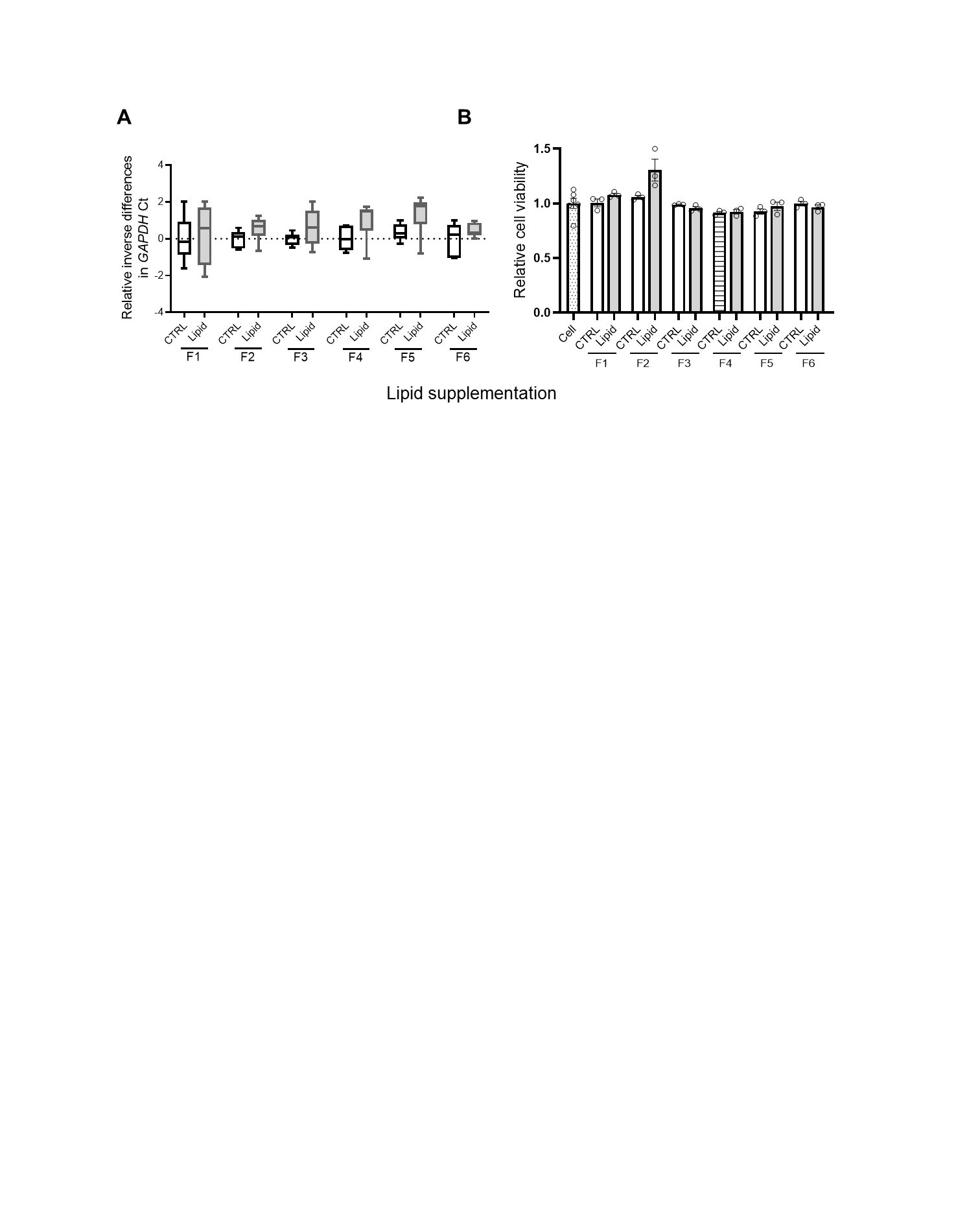

### Figure S6. Lipid fractions from mosquito EVs do not alter cell viability

(A, B) Huh7 cells were infected with DENV at MOI of 0.1 upon supplementation with 0.1 µl of mosquito EV-lipid fractions (F1-6). CTRL indicates supplementation with the corresponding solvents.

(A) Cell viability was evaluated by calculating relative inverse differences in *GAPDH* Ct values. Results are presented as Tukey box and whiskers derived from at least 6 biological replicates collected from multiple experiments.

(B) Cell viability was evaluated using CyQUANT NF Cell proliferation Assay at 72 hours post treatment. Bars show means ± SEM. Points represent biological repeats.

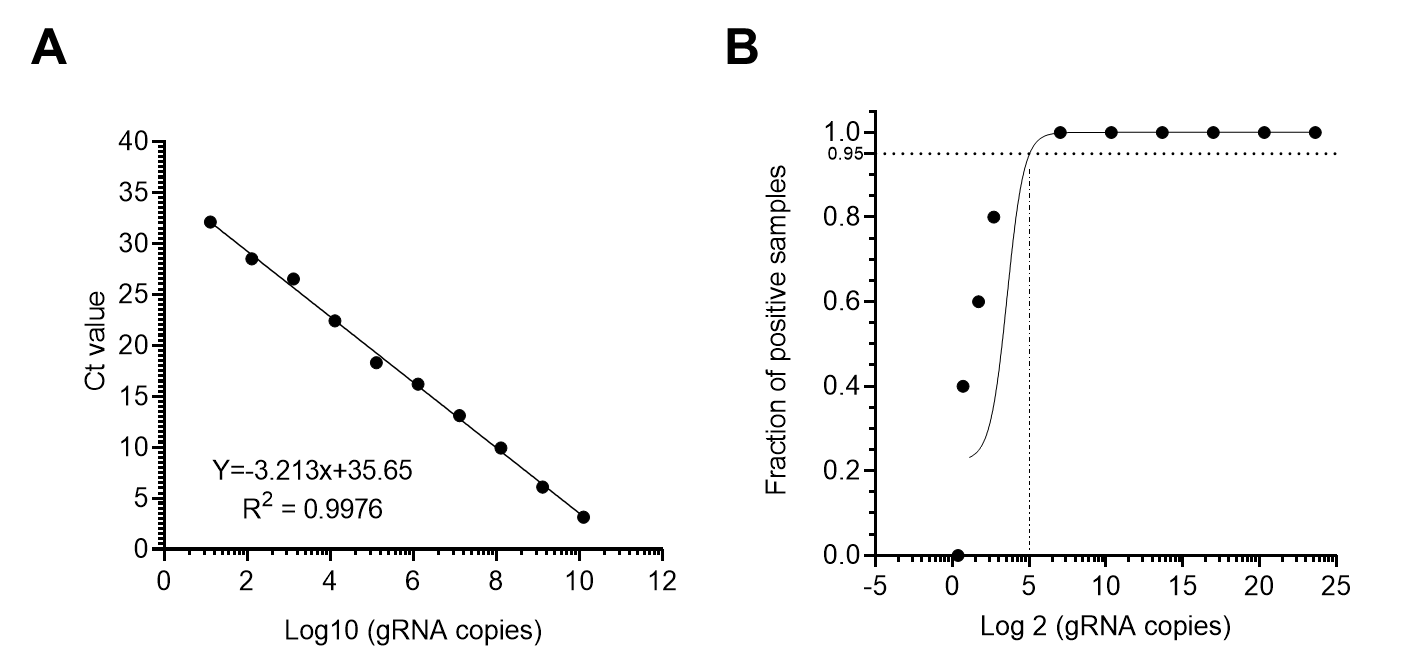

### Figure S7. Standard for absolute quantification of WNV gRNA

*In vitro* transcribed RT-qPCR target was serially diluted from 1.29 x 10^10^ to 1.29 copies before quantification by one-step RT-qPCR.

(A) Standard curve for absolute quantification of gRNA copies. Points represent the averaged Ct values calculated from 4 repeats.

(B) Limit of Detection (LoD) for WNV gRNA, determined at 95% confidence, is 25 copies (5²). The fractions of positive samples were computed based on 4 or 5 replicates.

### Table S1. Description of EV-protein extracts

Each ID indicates a different extraction event. Nd, not detected.

|  | **Details of the EV collection** | | | | **Details of the EV lysates used for extraction** | | | | | **Characteristics of the EV-protein extracts** | | | | | |
| --- | --- | --- | --- | --- | --- | --- | --- | --- | --- | --- | --- | --- | --- | --- | --- |
| **ID** | **Cell type** | **Number of flasks** | **Estimated total cell number (million)** | **Volume of EV lysate in RIPA (µl)** | **Volume (µl)** | **Estimated cell number (million)** | **Protein quantity (µg)** | **DNA quantity (µg)** | **RNA quantity (µg)** | **Volume (µl)** | **Estimated cell number of origin (cells/µl)** | **Protein quantity (µg)** | **Concentration of proteins (µg/µl)** | **DNA quantity (µg)** | **RNA quantity (µg)** |
| Prot 6.1 | Aag2 | 3 T175 | 45 | 800 | 56 | 3.15 | 112 | 0.056 | 1.12 | 50 | 63 000.00 | 81 | 1.62 | nd | nd |
| Prot 6.2 |  |  |  |  |  |  | 112 | 0.056 | 1.12 | 50 |  | 87.5 | 1.75 | nd | nd |

### Table S2. Description of EV-lipid extracts

Each ID indicates a different extraction event. Nd, not detected.

|  | **Details of the EV collection** | | | | **Details of the EV lysates used for extraction** | | | | | **Characteristics of the EV-lipid extracts** | | | | |
| --- | --- | --- | --- | --- | --- | --- | --- | --- | --- | --- | --- | --- | --- | --- |
| **ID** | **Cell type** | **Number of flasks** | **Estimated total cell number (million)** | **Volume of EV lysate in RIPA (µl)** | **Volume of lysate used for lipid extraction (µl)** | **Estimated cell number (million)** | **Protein quantity (µg)** | **DNA quantity (µg)** | **RNA quantity (µg)** | **Volume (µl)** | **Protein quantity (µg)** | **Concentration of proteins (µg/µl)** | **DNA quantity (µg)** | **RNA quantity (µg)** |
| LIP 1.1 | Aag2 | 2 T175 | 18 | 600 | 100 | 3 | 108 | 19 | 23 | 50 | 8.5 | 0.17 | 1.1 | 1 |
| LIP 2.2 |  | 6 T175 | 54 | 1500 | 90 | 3.24 | 110.7 | 14.8 | 29.6 | 50 | 26.5 | 0.53 | nd | nd |
| LIP 2.5_1 |  |  |  |  |  |  |  | 8.9 | 28.6 | 50 | 16.5 | 0.33 | nd | nd |
| LIP 2.5_2 |  |  |  |  |  |  |  | 9.55 | 26.2 | 50 | 18 | 0.36 | nd | nd |
| LIP 2.5_3 |  |  |  |  |  |  |  | 10.64 | 27.4 | 50 | 14 | 0.28 | nd | nd |
| LIP 6.1 |  | 3 T175 | 45 | 800 | 56 | 3.15 | 112 | 0.056 | 1.12 | 50 | 24 | 0.48 | nd | nd |
| LIP 6.2 |  |  |  |  |  |  | 112 | 0.056 | 1.12 | 50 | 27 | 0.54 | nd | nd |
| LIP 7.1 |  | 3 T175 | 39 | 800 | 60 | 2.925 | 112 | nd | nd | 50 | 25.5 | 0.51 | nd | nd |
| hLIP 1.1 | Huh7 | 3 T175 | 36 | 990 | 90 | 3.27 | 112.5 | 12 | 15.2 | 50 | 65 454.55 | 25 | 0.5 | nd |

### Table S3. Description of lipid extracts from DENV.

Each ID indicates a different extraction event. Nd, not detected. Na, not analysed.

| **Details of samples used for lipid extraction** | | | | | | | **Characteristics of the lipid extracts** | | | |
| --- | --- | --- | --- | --- | --- | --- | --- | --- | --- | --- |
| **ID** | **Cell type** | **Infectious status** | **Quantity of DENV gRNA (copies/µl)** | **Volume (µl)** | **Total DENV gRNA copies** | **Protein quantity (µg)** | **Volume (µl)** | **Protein quantity (µg)** | **DNA quantity (µg)** | **RNA quantity (µg)** |
| C6/36_Mock | C6/36 | Mock | nd | 54 | nd | na | 50 | 0.104 | nd | nd |
| C6/36_DENV2 | C6/36 | DENV2 | 4.34 x10^7 | 52 | 2.25 x 10^9 | 0.55 | 50 | 0.122 | nd | nd |
| LLC_Mock | LLC-MK2 | Mock | nd | 14 | nd | 0.272 | 50 | 0.136 | nd | nd |
| LLC_DENV2 | LLC-MK2 | DENV2 | 1.26 x 10^8 | 28 | 3.25 x 10^9 | 0.22 | 50 | 0.146 | nd | nd |

### Table S4. Clinical score for mice inoculated with WNV with mosquito EV-lipid extracts

| **Conditions** | **DENV, no EV-lipids** | | | | | | | **DENV + EV-lipids (µl)** | | | | | | | | | | | | | | **No DENV, no EV-lipids** | | | |
| --- | --- | --- | --- | --- | --- | --- | --- | --- | --- | --- | --- | --- | --- | --- | --- | --- | --- | --- | --- | --- | --- | --- | --- | --- | --- |
| **dpi** | **CTRL** | | | | | | | **0.1** | | | | | | | **1** | | | | | | | **No inf** | | | |
|  | M1 | M2 | M3 | M4 | M5 | M6 | M7 | M1 | M2 | M3 | M4 | M5 | M6 | M7 | M1 | M2 | M3 | M4 | M5 | M6 | M7 | M1 | M2 | M3 | M4 |
| 1 | 0 | 0 | 0 | 0 | 0 | 0 | 0 | 0 | 0 | 0 | 0 | 0 | 0 | 0 | 0 | 0 | 0 | 0 | 0 | 0 | 0 | 0 | 0 | 0 | 0 |
| 2 | 0 | 0 | 0 | 0 | 0 | 0 | 0 | 0 | 0 | 0 | 0 | 0 | 0 | 0 | 0 | 0 | 0 | 0 | 0 | 0 | 0 | 0 | 0 | 0 | 0 |
| 3 | 0 | 0 | 1 | 0 | 0 | 0 | 0 | 1 | 0 | 0 | 0 | 0 | 0 | 0 | 0 | 0 | 0 | 0 | 0 | 0 | 0 | 0 | 0 | 0 | 0 |
| 4 | 0 | 0 | 1 | 0 | 0 | 0 | 0 | 2 | 0 | 0 | 0 | 0 | 0 | 0 | 1 | 1 | 2 | 0 | 0 | 0 | 0 | 0 | 0 | 0 | 0 |
| 5 | 1 | 1 | 1 | 0 | 0 | 0 | 1 | 2 | 0 | 0 | 0 | 0 | 0 | 0 | 1 | 2 | 1 | 0 | 0 | 0 | 0 | 0 | 0 | 0 | 0 |
| 6 | 1 | 1 | 1 | 0 | 0 | 0 | 1 | 4 | 1 | 1 | 0 | 0 | 0 | 1 | 4 | 4 | 1 | 0 | 0 | 0 | 0 | 0 | 0 | 0 | 0 |
| 7 | 0 | 0 | 0 | 1 | 0 | 0 | 1 | 5 | 0 | 1 | 1 | 0 | censor | 2 | 4 | 4 | 1 | 0 | 0 | 1 | 1 | 0 | 0 | 0 | 0 |
| 8 | 0 | 1 | 0 | 1 | 0 | 1 | 2 | 5 | 3 | 1 | 1 | 1 |  | 3 | 5 | 4 | 1 | 2 | 2 | 2 | 2 | 0 | 0 | 0 | 0 |
| 9 | 0 | 0 | 0 | 2 | 0 | 2 | 5 | 5 | 4 | 1 | 3 | 1 |  | 4 | 5 | 4 | 1 | 4 | 5 | 3 | 4 | 0 | 0 | 0 | 0 |
| 10 | 1 | 0 | 0 | 3 | 0 | 3 | 5 | 5 | 3 | 1 | 4 | 2 |  | 5 | 5 | 2 | 1 | 4 | 5 | 3 | 4 | 0 | 0 | 0 | 0 |
| 11 | 0 | 0 | 1 | 3 | 0 | 4 | 5 | 5 | 2 | 0 | 3 | 2 |  | 5 | 5 | 1 | 0 | 5 | 5 | 4 | 5 | 0 | 0 | 0 | 0 |
| 12 | 0 | 0 | 1 | 2 | 0 | 3 | 5 | 5 | 2 | 0 | 3 | 2 |  | 5 | 5 | 1 | 0 | 5 | 5 | 4 | 5 | 0 | 0 | 0 | 0 |

M, mouse; dpi, day post-infection, Died, Mouse deceased during blood collect

| Clinical score | Interpretation |
| --- | --- |
| 0 | Healthy mouse (baseline) |
| 1 | Weight loss, ruffled fur, lethargy, hunched posture, no paresis, normal gait |
| 2 | Weight loss (+), altered gait, breathing difficulty, limited movement in 1 hind limb |
| 3 | Emaciation, lack of movement, labored breathing, paralysis in 1 or both hind limbs |
| 4 | Moribund, cachexia |
| 5 | Dead |

### Table S5. List of primers.

| **Organism** | **Target** | **Name** | **Sequences (5’-3’)** | **References** |
| --- | --- | --- | --- | --- |
| DENV2 | NS5 D2 (+)gRNA | F | CTCCCTGAGTGGAGTGGAAG | (Yeh et al., 2023) |
|  |  | R | ACACGCACCACCTTGTTTTG |  |
|  | To produce DENV standard | F-T7-D2 | TAATACGACTCACTATAGGGCTCCCTGAGTGGAGTGGAAG |  |
|  | DENV (-) gRNA | F | CCTTTTCCAAATAATCCGCATC | (Vial et al., 2020) |
|  |  | R | ACCAACACAACAACAGCATC |  |
|  |  | Probe | HEX-CCTGTTCTTCATTTAGGCTAGGTTCTCCTTGT- BHQ-1 |  |
| WNV | Envelope protein E of strain IS98 | F | ATTCGGGAGGAGACGTGGTA | This study |
|  |  | R | CAGCCGCCAACATCAACAAA |  |
|  | To produce WNV standard | F-T7-WNV | TAATACGACTCACTATAGGGATTCGGGAGGAGACGTGGTA |  |
| ZIKV 835, ZIKV 911c | prM | F | TTGGTCATGATACTGCTGATTGC | (Lanciotti et al., 2008) |
|  |  | R | CCTTCCACAAAGTCCCTATTGC |  |
| CHIKV | Enveloppe glyceoprotein E1 | F | AAGCTYCGCGTCCTTTACCAAG | (Pastorino et al., 2005) |
|  |  | R | CCAAATTGTCCYGGTCTTCCT |  |
|  |  | Probe | FAM-CCAATGTCYTCMGCCTGGACACCTTT-TAMRA |  |
| *Mycoplasma* spp*.* | 16S rRNA | F | GGCGAATGGGTGAGTAACACG | (Sreedharan et al., 2017) |
|  |  | R | CGGATAACGCTTGCGACTATG |  |
| Human | GAPDH | F | AAATCAAGTGGGGCGATGCT | (Koh et al., 2006; Tang et al., 2015) |
|  |  | R | AAGCAGTTGGTGGTGCAGGA |  |
|  | RPL13A | F | AACAGCTCATGAGGCTACGG | Maarifi et al 2021. EMBO J |
|  |  | R | TGGGTCTTGAGGACCTCTGT |  |
|  | ACTB | F | CTGGAACGGTGAAGGTGACA |  |
|  |  | R | AAGGGACTTCCTGTAACAATGCA |  |
|  | B2M | F | TGCTGTCTCCATGTTTGATGTATCT |  |
|  |  | R | TCTCTGCTCCCCACCTCTAAGT |  |
|  | GAPDH | F | TGCACCACCAACTGCTTAGC |  |
|  |  | R | GGCATGGACTGTGGTCATGAG |  |
|  | IFN-β | F | TGCTCTCCTGTTGTGCTTCTC |  |
|  |  | R | CAAGCCTCCCATTCAATTGCC |  |
|  | CXCL10 | F | CGCTGTACCTGCATCAGCAT |  |
|  |  | R | GCAATGATCTCAACACGTGGAC |  |
|  | Mx1 | F | AAGCTGATCCGCCTCCACTT |  |
|  |  | R | TGCAATGCACCCCTGTATACC |  |
|  | Serl1 L | Sel1L_F | TGGGTTTTCTCTCTCTCCTCTG | Sun et al 2014. PNAS |
|  |  | Sel1L_R | CCTTTGTTCCGGTTACTTCTTG |  |
|  | Derlin1 | Derlin-1_F | CACCTCAGTTTTTGTACCGCTG | Fan et 2020 |
|  |  | Derlin-1_R | TCACTGGTCTCCAAGTCGAAAG |  |
|  | Edem1 | EDEM1-F | CGGACGAGTACGAGAAGCG | Li et al; 2021. Aging |
|  |  | EDEM1-R | CGTAGCCAAAGACGAACATGC |  |
|  | Herpud1 | Herpud1-F | ACTCCTCGCTGAGCAGATTT | Ho adn Chan, 2015. FEBS letters |
|  |  | Herpud1-R | CTCTGTCTGAACGGAAACCA |  |
|  | Hrd1 | Hrd1-F | CCAGTACCTCACCGTGCTG | Hu et al, 2014. Genes & Dev |
|  |  | Hrd1-R | GCCTCTGAGCTAGGGATGC |  |

### Dataset S1. Composition of mosquito EV-lipids

Quantitative targeted lipidomics on mosquito EV-lipid extracts. Data derived from three biological replicates (rep1- 3). Quantity for PL, SM, NL, and FA. Last worksheet shows summary of quantification per lipid type. Nd, not detected.

### Dataset S2. Composition of SPE fraction from mosquito EV-lipids.

SPE fractions 1-6 were subjected to untargeted lipidomics. Normalized peak area for all identified metabolites is presented. Other worksheets aggregate the results per lipid class (i.e., Cer, SM, PS, PE, PC, PL precursor, TAG, DAG, FA and NEA) and show calculated within-class proportion, %.

Cer, Ceramide; SM, Sphingomyelin; PC, Phosphatidylcholine; PS, Phosphatidylserine; PE, Phosphatidylethanolamine; PI, Phosphatidylinositol; TAG, triacylglycerides; DAG, diacylglycerides; MAG, Monoacylglycerides; NAE, N-acylethanolamine; ST, sterol.

### Dataset S3. Effect of mosquito EV-lipids and DENV on Huh7 lipidome

Huh7 cells were inoculated with 0.1 µl of mosquito EV-lipids, DENV at MOI 0.1 or both lipids and DENV. Control cells were neither supplemented with lipids nor infected. Untargeted lipidomics of cells was conducted after 4 h. Tables show metabolite names, MS peak surface in three replicates for each condition, and statistics [f.value, p-value, -log10(p)].

Grey cells indicate metabolites not significantly regulated (|log2 fold change| < 1; p-value > 0.05).

Cer, Ceramide; SM, Sphingomyelin; PC, Phosphatidylcholine; PS, Phosphatidylserine; PE, Phosphatidylethanolamine; PI, Phosphatidylinositol; TAG, triacylglycerides; DAG, diacylglycerides; MAG, Monoacylglycerides; NAE, N-acylethanolamine.
